# Inhibition of adenylyl cyclase 1 (AC1) and exchange protein directly activated by cAMP (EPAC) restores ATP-sensitive potassium (K_ATP_) channel activity after chronic opioid exposure

**DOI:** 10.1101/2025.02.03.636278

**Authors:** Amanda H. Klein, Sabbir Alam, Kayla Johnson, Christian Kriner, Brie Beck, Bethany Nelson, Cassidy Hill, Belle Meyer, Jonas Mellang, Val J. Watts

**Affiliations:** Department of Pharmacology and Toxicology, University at Buffalo, Buffalo, NY; Department of Pharmacy Practice and Pharmaceutical Sciences, University of Minnesota, Duluth, MN; Borch Department of Medicinal Chemistry and Molecular Pharmacology, Purdue University, West Lafayette IN; Purdue Institutes for Integrative Neuroscience (PIIN), Drug Discovery (PIDD), Cancer Research (PICR), and Inflammation, Immunology and Infectious Disease (PI4D), Purdue University, West Lafayette IN

**Keywords:** Adenylyl Cyclase, cAMP Signaling, EPAC, opioid, tolerance, withdrawal, potassium channel

## Abstract

Prolonged exposure to Gαi/o receptor agonists such as opioids can lead to a sensitization of adenylyl cyclases (ACs), resulting in heterologous sensitization or cyclic AMP (cAMP) overshoot. The molecular consequences of cAMP overshoot are not well understood, but this adaptive response is suggested to play a critical role in the development of opioid tolerance and withdrawal. We found that genetic reduction of AC1 and simultaneous upregulation of ATP-sensitive potassium (K_ATP_) channel subunits, SUR1 or Kir6.2, significantly attenuated morphine tolerance and reduced naloxone-precipitated withdrawal. *In vitro* models utilized an EPAC2-GFP-cAMP biosensor to investigate sensitization of adenylyl cyclase in SH-SY5Y neuroblastoma cells and HEKΔAC3/6 knockout cells. Acute application of DAMGO significantly decreased the cAMP signal from the EPAC2-GFP-cAMP biosensor, while chronic DAMGO administration resulted in enhanced cAMP production following AC stimulation. Inhibition of cAMP overshoot was observed with naloxone (NAL), pertussis toxin (PTX), and the neddylation inhibitor, MLN4924 (Pevonedistat), as well as co-expression of β-adrenergic receptor kinase C-terminus (βARK-CT). After establishment of the AC1-EPAC sensitization in the *in vitro* models, we found that inhibition of AC1 or EPAC enhanced potassium channel activity after chronic morphine treatment, using a thallium-based assay in SH-SY5Y cells. Similar data were obtained in mouse dorsal root ganglia (DRG) after chronic morphine treatment. This study presents evidence for investigating further AC1 signaling as a target for opioid tolerance and withdrawal, by increasing EPAC activity and affecting potassium channels downstream of opioid receptors.

**Graphical Abstract.**
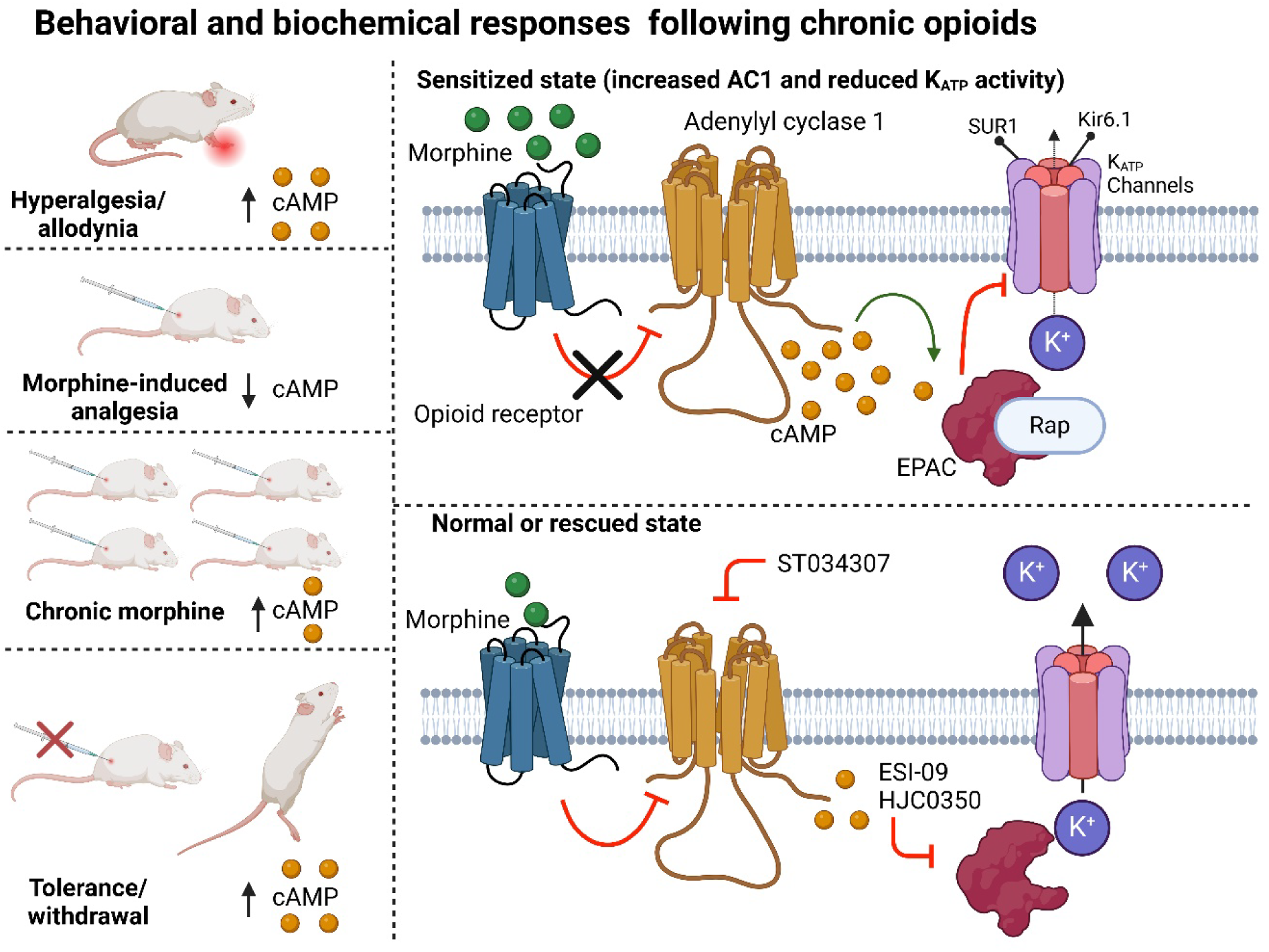

## 1. Introduction

Among the ten known AC isoforms, adenylyl cyclase 1 (AC1) is highly expressed in the central nervous system (CNS) [1] and is critically involved in pain signaling and nociceptive pathways, influencing synaptic plasticity and the nervous system’s response to painful stimuli [2]. AC1 is robustly activated by Ca^2+^/calmodulin (CaM), forskolin, and Gαs, and inhibited by Gαi and Gβγ subunits[3]. AC1 is also one of several AC isoforms that undergo heterologous sensitization or cAMP overshoot after chronic activation of Gαi/o-coupled receptors, such as mu opioid receptors [4]. Increased AC activity has been demonstrated using several *in vitro* models of chronic opioid exposure [5, 6]. When morphine is withdrawn, the morphine-induced inhibition of AC is eliminated, uncovering a phenomenon known as heterologous sensitization, AC supersensitization, or cAMP overshoot [7, 8]. These observations, along with subsequent research, pointed to a potential mechanism linking heterologous sensitization of cAMP to opioid tolerance and dependence, and potentially withdrawal [7, 9]. The precise molecular mechanisms behind cAMP overshoot remain poorly understood, and more importantly, the physiological impacts of elevated cAMP on downstream signaling pathways are also underdeveloped.

cAMP has several direct actions on neurons within the peripheral and/or central nervous system, which can increase synaptic transmission and/or change hyperpolarization-activated currents (Ih) [10]. Another target specifically of cAMP is EPAC (exchange protein directly activated by cAMP). EPAC1 and EPAC2, found within the CNS, are guanine nucleotide exchange factors for Rap1, part of the Ras family of G-proteins, contributing to the effective signaling of GPCRs. EPAC expression is increased in morphine tolerant rats, and intrathecal administration of the inhibitor ESI-09 reduces morphine tolerance [11]. The mechanisms of how EPAC facilitates hyperalgesia, are still being understood [12], but potential mechanisms could involve phospholipase activity in non-peptidergic C-fibers [13], or through modulation of ion channels[14]. cAMP-mediated activation of EPAC inhibits ATP-sensitive potassium (K_ATP_) channels via a Ca^2+^-dependent mechanism, indicating that under certain conditions cAMP conveys inhibitory information to potassium channels[15]. Since K_ATP_ channels are involved in downstream signaling of opioid receptors [16], and their expression and function are correlated with hypersensitivity in pain conditions [17, 18], we wanted to determine if modulation of expression and function of AC and K_ATP_ channels could impact the development of morphine tolerance and withdrawal in mice. We also wanted to determine if *in vitro* cellular models of chronic opioid administration would reflect enhanced cAMP activity using an EPAC2-GFP cAMP biosensor (cADDis; cAMP Difference Detector *in situ*). Finally, we wanted to explore if inhibition of EPAC activity after chronic opioid exposure would rescue endogenous K_ATP_ channel activity in a human cell line and mouse dorsal root ganglion (DRG). These results demonstrate that heterologous sensitization of AC and cAMP activity increases EPAC2 and possibly contributes to nociceptor hyperactivity after chronic opioid exposure. Future translational studies should consider AC1 and/or EPAC as a potential therapeutic target for opioid tolerance and withdrawal, in addition to chronic pain.

## 2. Materials and methods

### 2.1 Animals

Adult male and female C57Bl6/6NCRL mice were purchased from Charles River Laboratories (Raleigh, NC) at six weeks of age. The animal experiment protocols were approved by the Institutional Animal Care and Use Committees of the University of Minnesota and the University at Buffalo. Approximately equal numbers of male and female mice were used in every experiment. Male and female animals were separated by sex on the day of experimentation and equipment was thoroughly cleaned between each group of animals. Animals were randomized to the treatment group(s), and the experimenter was blinded to the treatment group, and unblinded after data collection and analysis was completed.

### 2.2 Mouse Virus administration

Ten microliters of each virus (Vector Biolabs, Malvern, PA, United States) was delivered by direct lumbar puncture in awake mice[19]. Behavioral assessments began 4 weeks post AAV9- virus injection and ∼1 week post Ad-virus. Groups tested were as follows: AAV9-RFP-U6-m- Adcy1-shRNA (shAAV-252120) + Ad-m-Abcc8 (ADV-251792), AAV9-RFP-U6-scrmb-shRNA (7045)+ Ad-m-Abcc8, AAV9-RFP-U6-m-Adcy1-shRNA + Ad-CMV-Null (1300), AAV9-RFP- U6-m-Adcy1-shRNA + Ad-m-Kcnj11(ADV-262651), AAV9-RFP-U6-scrmb-shRNA + Ad-m-Kcnj11, and AAV9-RFP-U6-scrmb-shRNA + Ad-CMV-Null. Several rounds of injections in separate cohorts of mice were performed to counterbalance sex and virus combination, control viruses were run with every group of animals.

### 2.3 Behavior

All behavioral experiments were performed in a blinded fashion. Mice were acclimated to plexiglas boxes on an elevated mesh platform at least three times before the start of behavioral testing. One week after Ad administration, a cumulative morphine dose response was performed by injection of 1 mg/kg, 2 mg/kg, 5 mg/kg, 10 mg/kg and 15 mg/kg morphine (sc, in saline) every thirty minutes and subsequent mechanical paw withdrawal measurements. Morphine tolerance was assessed by injecting morphine (15 mg/kg s.c.) twice per day for 5 days. Mechanical paw withdrawal measurements were again taken before and 30 min after morphine administration to measure opioid induced hyperalgesia and opioid tolerance, respectively. One day after completion of morphine tolerance, a second morphine dose response curve was performed. Precipitated withdrawal was performed three days after morphine tolerance testing. Thirty minutes after systemic morphine (15 mg/kg, sc, in saline), mice were placed in clear acrylic chambers with three-way mirrors and behaviors were recorded by video camera for off-line analysis for 10 min. Three hours post morphine injection, a dose of 1 mg/kg naloxone (ip in saline) was administered, and mice were recorded again. Behavioral scoring of withdrawal was done by recording the number of jumps (defined as the simultaneous removal of all four paws from the ground) and number of times the animal reared (standing on the hind legs to explore the environment)[20].

### 2.4 Quantitative PCR (qPCR)

Lumbar spinal cord and DRG from mice were collected after completion of behavioral experiments to determine knockdown or upregulation of AC1 and SUR1/Kir6.2 subunits, respectively. Tissues were flash frozen in liquid nitrogen and stored at −80oC until RNA processing using Direct-zol RNA Miniprep Kits (Zymo Research, Irvine, CA) with TRI Reagent™ Solution (Thermo Fisher Scientific). cDNA was synthesized using random nonameric primers (Integrated DNA Technologies, Coralville, IA) from amounts of RNA that were standardized for each experiment using the Omniscript RT Kit (Qiagen) or the High-Capacity cDNA Reverse Transcription Kit (Thermo Fisher Scientific). cDNA (2µl of 200µl cDNA reaction mix) was taken for qPCR carried out using either Sso Advanced Universal SYBR Green Supermix (Biorad Laboratories, Hercules, CA) on a BioRad CFX Real-Time System C100 Touch Thermal Cycler or LightCycler 480 SYBR Green I Master mix (Roche Diagnostics Corporation, Indianapolis, IN) on a LightCycler 480 Instrument II. Specific primers were used to quantify the cDNA levels of various housekeeping genes (Gapdh, 18S rRNA) and Abcc8, Abcc9, Kcnj11, Kcnj8, Adcy1, and Adcy8 (Supplemental Table 1)[21]. cDNA copy numbers were quantified by the C_T_ values as determined by the software. Samples were run in triplicate against a ≥5 point, 10-fold serial dilution of gene specific cDNA standard. The temperature and cycling protocol were typically as follows: 95 °C for 10 sec, 60 °C for 10 sec and 72 °C for 20 sec and was repeated for 40 cycles. Negative RT-PCR controls were performed to assess the presence of genomic DNA.

SH-SY5Y cells were obtained from and maintained through continuous culture (American Type Culture Collection, Manassas, VA, CRL-2266). SH-SY5Y cells were used for qPCR analyses of ADCY1, ADCY3, ADCY5, ADCY8, RAPGEF3, and RAPGEF4 expression after morphine exposure (Supplemental Table 2). In 6-well plates cells were seeded equally and treated with 0, 5, 10, 25, 50, or 100 uM morphine for 72 hrs in quadruplicate. Cells were collected and processed for mRNA quantification as stated above. Morphine concentrations were based on BCA and MTT assays which determined that morphine concentrations >100uM may be toxic (data not shown). qPCR data were normalized to housekeeping genes (18S rRNA, GAPDH).

### 2.5 Cells and reagents: cADDis cAMP Assay

HEK-ACΔ3/6 cells stably expressing human AC1 was generated by knocking out AC3 and AC6 with CRISPR/Cas9 and transfecting with AC1 expression vector, followed by selection for stable AC1 cells with G418 [22]. Dulbecco’s modified Eagle’s medium (DMEM) (Catalog # 11995065 Gibco; 10-090-CV, Corning), penicillin-streptomycin-amphotericin B solution (antibiotic-antimycotic), cell dissociation buffer TrypLE (Catalog # 12605028), and OptiMEM (Catalog # 31985070) were purchased from ThermoFisher Scientific, USA. Bovine calf serum, and Fetal Clone I were obtained from Hyclone (GE Healthcare, Pittsburgh, PA). The assay plates were coated with poly-D lysine (PDL, Sigma Cat# P6407, 200 ng/mL) for 2 hours at room temperature, washed 6 times with ddH_2_O, and then dried overnight before use. We utilized the cADDis, Epac2-GFP cAMP biosensor, a genetically encoded fluorescent biosensor, to monitor cellular cAMP levels in real-time. This sensor employs a modified version of green fluorescent protein (GFP) called mNeoGFP, which is positioned within the EPAC2 guanine nucleotide exchange factor. The cAMP-sensor changes fluorescence in response to cAMP binding and unbinding, with upward and downward shifts, and we specifically used the green upward version (Montana Molecular; Catalog #U0200G) to monitor intracellular cAMP level changes [23].

### 2.6 Viral transduction of SH-SY5Y and HEK cells

SH-SY5Y cells or HEK-ACΔ3/6 cells stably expressing AC1 (HEK AC1 cells) were cultured in DMEM, with 10% fetal bovine serum (FBS) and penicillin-streptomycin at 37°C in 5% CO_2_. For BacMam transduction, the cells were resuspended in DMEM to a density of 70,000 cells per well, supplemented with 2 mM sodium butyrate (Sigma, St. Louis, MO). Each well received 5 × 10^8 viral genomes of the cAMP sensor in Bacmam, with the viral load carrying the μ-opioid receptor (MOR) and AC1 at 6 x 10^7 and 4 x 10^7 viral genomes per well, respectively. The cell and transduction mixture were then plated into 96-well plates and incubated for 24 to 48 hours at 37°C in a 5% CO_2_ atmosphere. Before fluorescence plate reading and/or Cytation 3 imaging, the media was replaced with Dulbecco’s phosphate-buffered saline (DPBS) supplemented with 0.9 mM Ca^2+^ and 0.5 mM Mg^2+^.

### 2.7 cAMP signaling in HEK and SH-SY5Y cells

To investigate cAMP signaling in SH-SY5Y neuroblastoma cells and HEK AC1 cells, we adapted the protocol from Xia and colleagues (2011) [24]. The cells were transduced with the cAMP biosensor in the absence or presence of indicated viruses (AC1, MOR, and/or βARK-CT) for 24 hours and then acute and prolonged drug treatments were carried out as described. For mechanistic overshoot experiments, cells were pretreated with pertussis toxin (Ptx, 25 ng/mL) for 2 hours or MLN4924 (1 μM) or naloxone (1 μM) for 30 minutes as indicated. Following the pretreatment/transduction, the cells were incubated with DAMGO (1 μM) in combination with Ptx or MLN4924, or control for 18 hours. cAMP stimulation was carried out using 5 μM A23187 in the presence of naloxone to prevent residual DAMGO-mediated inhibition of cAMP. The stimulation period was followed by maximal stimulation with forskolin (FSK), naloxone, and the phosphodiesterase inhibitor, isobutylmethylxanthine (IBMX). For studying exogenous cAMP overshoot in SH-SY5Y neuroblastoma cells, the cells were transduced with cAMP-sensor, MOR, and AC1 for 24 hours and experiments carried out as described above. Prior to each experiment, cell health was assessed under a brightfield microscope.

### 2.8 cADDis Fluorescence Imaging

To capture a time-lapse series of dynamic cAMP activity, the SH-SY5Y cells were incubated with DPBS supplemented with Ca²⁺ and Mg²⁺ for 30 minutes. For cADDis time-lapse fluorescence imaging, images were acquired using an inverted Cytation 3 microscope fitted with a 20×0.9 NA objective and laser autofocus was used to automatically adjust focus. Images were acquired at intervals of 1 minute and 45 seconds. Fluorescence plate reader experiments were conducted using the BioTek Synergy Neo2 plate reader with 96-well plates. Fluorescence was detected (with 45 sec/read) using a 485/20 nm excitation wavelength and a 520/20 nm emission filter, with optics positioned at the bottom. A gain of 100 was applied, and the light source was a Xenon flash with low lamp energy. The standard dynamic range was used, along with a normal read speed and a delay of 0 msec. Measurements were taken with 10 data points, and the read height was set to 4.5 mm.

### 2.5 Potassium flux

Potassium channel activity was determined by thallium (Tl⁺) flux assay (Brilliant Thallium Assay Express Kit, Ion Biosciences, San Marcos, TX, 11000- 100). Although described in great detail previously[25], our methods closely follow the instructions provided by the vendor. Briefly, cells were placed on PDL (P7280, Sigma Aldrich, St. Louis, MO) coated glass bottom 35mm dishes (Cellvis, D35-14-1.5N, Sunnyvale, CA). SH-SY5Y cells were allowed to be cultured for three days with or without morphine (Spectrum Chemical, New Brunswick NJ) or DAMGO before imaging. Dishes treated with morphine for 72 hours had the media changed every day. On the day of imaging, cells were loaded with Brilliant Thallium Reagent for one hour with or without Glyburide (1uM), ESI-09 (5uM), or ST034307 (30uM). Stock solutions were initially made in DMSO and diluted to a final concentration of less than 5% DMSO. Cells were imaged using excitation and emission filters at 395/509nm with a Nikon Ti2-E inverted research microscope with X-Light V2 LFOV spinning disk confocal system or a Leica DMi 8 inverted microscope with a TCS SP8 confocal microscope system. Images were taken every 2 seconds for at least 30 seconds before and five minutes after the addition of thallium solution, with or without 100uM diazoxide, delivered by perfusion tubing into the illuminated area.

### 2.10 Mouse DRG Culture and Labeling

DRG were removed from the spinal column of adult male and female mice (>6 weeks age) under a dissection microscope as described previously [26, 27]. Briefly, DRG were minced with fine spring scissors and incubated with papain (LS003126, Worthington Biochemical, Lakewood NJ) and then a collagenase (LS004176, Worthington Biochemical, Lakewood NJ) solution in HBSS for ten minutes at 37°C. DRG were triturated with EMEM media (#10-010-CV, Corning Manassas, VA) supplemented with vitamins and penicillin streptomycin (#15140-122, Gibco). Cells were cultured overnight on PDL coated glass bottom dishes. The next morning, cells were loaded with markers for isolectin B4 (IB4, #132450, Invitrogen) and Cholera B Toxin (ChBTx, #C22843, Invitrogen) for one hour before loading with thallium indicating dye (see above). Images were additionally acquired for detection of IB4 and ChBTx at Cy5 (647/670nm) and TRITC (555/565 nm) related wavelengths. Cells positive for IB4 are thought to be non-peptidergic and express P2X3 and GDNF receptors, while cells positive for ChBTx are myelinated fibers [28]. Cells without labels for either IB4 or ChBTx are thought to be peptidergic C-fibers and express trkA receptors. Very few (<10%) cells express markers for both IB4 and ChBTx, and are presumed A-delta fibers[29].

### 2.5 Data Analysis and statistics

Behavioral data are reported as raw values without normalization and qPCR data were log transformed to meet the requirements of normality for ANOVA. Fluorescence measurements from plate reader time-lapse imaging were analyzed using Fiji ImageJ analysis software[30]. The analysis included kinetic frame alignment and defining regions of interest (ROIs) to extract average traces using the multi-ROI tool. Peak response area was determined using five consecutive data points at the peak, while total response was assessed by calculating the total area of the response after respective drug treatment. The area under the curve (AUC) was calculated using GraphPad Prism with the trapezoidal method, summing the areas of trapezoids formed between successive data points and treating them as rectangles. Potassium flux assays were analyzed using Nikon Elements Software and/or Image J. Baseline fluorescence measurements were averaged over 30 seconds and the maximum percent change from baseline ([(peak response/baseline average)-baseline)] x 100) and the area under the curve (AUC) was calculated based on the trapezoidal rule two minutes after the addition of thallium.

Data are presented by either mean ± SEM or median ± 95% Confidence Intervals (CI) for parametric and non-parametric data, respectively. Data analysis and statistical significance tests were performed using GraphPad Prism 9 or 10. Non-parametric comparisons were performed by Kruskal–Wallis test with Dunn’s multi-comparisons. Parametric statistics were performed with one-way or two-way ANOVA with Tukey’s test applied for multiple comparisons across all groups or Dunnett’s test against a control group. Pairwise comparisons were conducted through two-tailed unpaired t-tests. Significance levels were represented as follows: “ns” for p > 0.05, “*” for p ≤ 0.05, “**” for p ≤ 0.01, “***” for p ≤ 0.001, and “****” for p ≤ 0.0001.

## 3. Results

### 3.1 Upregulation of K_ATP_ channel subunits and downregulation of AC1 reduces morphine tolerance and withdrawal symptoms in mice

We previously demonstrated in mice that inhibition of AC1, either pharmacologically or by viral knockdown (AAV9-shRNA), was able to decrease tolerance to morphine [31]. To determine if the reduction in Adcy1 or the upregulation of neuronal K_ATP_ channel subunits, Abcc8 or Kcnj11, have further impacts on the development of morphine tolerance or withdrawal, we utilized a viral strategy. Mice injected with Adcy1-shRNA had significantly higher paw withdrawal thresholds compared to control vector mice, especially when combined with Ad-Abcc8 (Figure 1A) or Ad- Kcnj11 (Figure 1B). The impact of loss of Adcy1 or upregulation of Abcc8/Kcnj11 was smaller for pre-morphine paw withdrawal measurements (Figure 1C) than post-morphine measurements (Figure 1D). After morphine tolerance, mice underwent precipitated withdrawal by naloxone injection. The number of recorded jumps was significantly decreased by loss of Adcy1 or upregulation of Abcc8/Kcnj11 (Figure 1E), but rearing was not affected (Figure 1F). Dose response curves of morphine were performed before tolerance and after tolerance was established. The thresholds pre-vs-post tolerance were shifted at least 1 gram for the control vector-treated mice but were least affected in the Adcy1-shRNA+Ad-Abcc8 and Adcy1-shRNA+Ad-Kcnj11 treated mice (Figure 1G).

**Figure 1.**
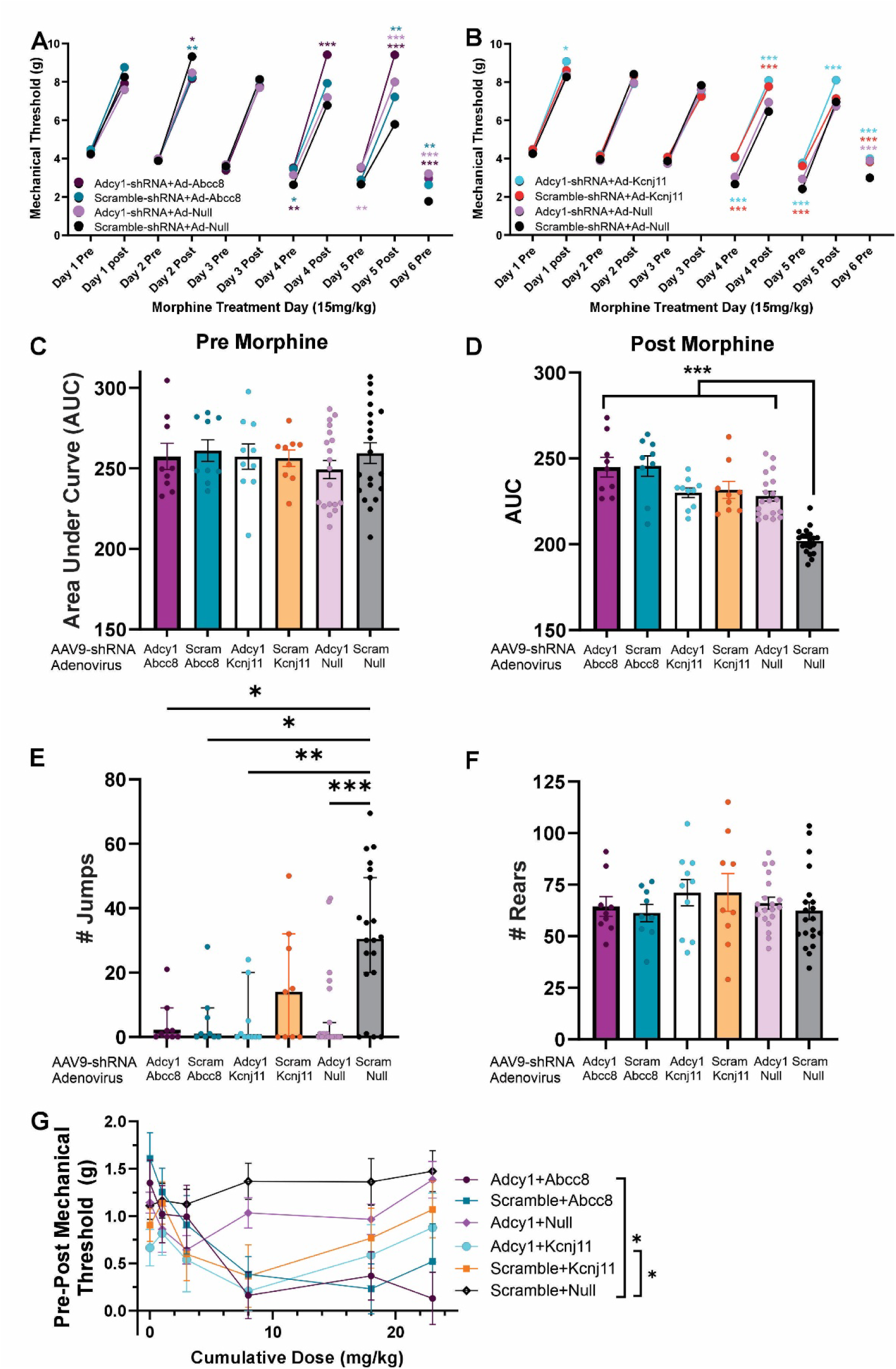
Upregulation of K_ATP_ channel subunits and downregulation of AC1 reduce morphine tolerance and withdrawal in mice. (A) Intrathecal administration of AAV9-Adcy1-shRNA to knock down AC1 expression increased mechanical paw withdrawal thresholds compared to control associated adenovirus (AAV9-Scrambl-shRNA). Mice treated with morphine (15mg/kg, twice daily, 5 days). Upregulation of SUR1 by an adenovirus (Ad-Abcc8) treated animals have attenuated morphine tolerance compared to mice treated with a control adenovirus (Ad-Null). Adcy1+Abcc8 (Adcy1-shRNA + Ad-Abcc8) treated mice compared to Scramble-Null (control) animals (two-way ANOVA w/ Dunnett’s post hoc) (B) Upregulation of Kir6.2 by an adenovirus (Ad-Kcnj11) and Adcy1-shRNA treatment also attenuated morphine tolerance compared to mice treated with a control AAV (Scram) and adenovirus (Null). (C) Cumulative dose response testing was conducted prior to morphine tolerance and (D) on the day after morphine tolerance (1, 2.5, 5, 10, 15 mg/kg). The area under the curve was calculated compared to AAV and Ad control treated mice (Scramble-Null) (one-way ANOVA w/ Dunnett’s post hoc). (E) The day after morphine tolerance was completed (Day 6), Mice were given an injection of morphine (15mg/kg), followed by naloxone and the number of jumps was recorded. Jumping was greatly increased for Adcy1-Abcc8 and Adcy1-Kcnj11 treated mice compared to control virus (Scramble-Null, Kruskal-Wallis, with Dunn’s post-hoc). (F) Rearing behaviors were also recorded in these animals at the same time by blinded reviewers. (G) Dose response curves for morphine antinociception were taken before morphine tolerance and after 5 days of chronic morphine exposure. The thresholds of morphine antinociception were most greatly increased at the higher cumulative doses of morphine for Adcy1-Abcc8 and Adcy1-Kcnj11 treated mice compared to control virus (Scramble-Null, two-way ANOVA with Dunnett’s post-hoc). Data presented as mean ± SEM with n=9-10 per group with male and female animal data combined. Data in E presented as median ± 95% CI. Statistical significance is indicated as follows: *p < 0.05, **p < 0.01, ***p < 0.001, ****p<0.0001.

### 3.2 Adenylyl cyclase activity in HEKΔAC3/6 KO cells expressing exogenous AC1 and MOR using the cADDis biosensor

The cADDis (i.e. EPAC2-GFP cAMP biosensor) contains the cAMP binding domain of EPAC2 fused to a circularly permuted form of mNeonGreen where increases in cAMP lead to increases of fluorescence. The presence of the EPAC2 binding domain also allows us to investigate the impact of acute and chronic opioid exposure on EPAC signaling *in vitro* [32]. Initial studies sought to validate the utility of cADDis biosensor in a previously characterized recombinant system. Specifically, we expressed the EPAC2-GFP cAMP-sensor in HEKΔAC3/6 KO cells that stably express AC1 (HEK-AC1 cells) and transiently express the µ-opioid receptor (MOR)[33]. cAMP activity was stimulated using the calcium ionophore A23187, which specifically activates recombinant AC1 activity in this model [33]. The expression and activity of the EPAC2-GFP cAMP sensor were confirmed through GFP images, demonstrating its intracellular localization and expression (Figure 2A, B). Figure 2C shows representative time-lapse captures before and after the addition of A23187 to selectively activate AC1. The subsequent addition of DAMGO reduced the A23187-stimulated fluorescent response as shown. The dynamic nature of the cADDis was further revealed following the final addition of forskolin (10 μM) in the presence of IBMX to maximally stimulate cAMP accumulation in the absence of enzymatic degradation (Fig. 2C). Having established this recombinant model, quantitative studies were then carried out. These studies revealed a robust increase in fluorescence and cAMP kinetics upon the addition of the non-specific AC activator, forskolin (Figure 2D-F) or A23187 (Figure 2G-I). The initial stimulation with either forskolin or A23187 reached a plateau within 10 minutes, with the signal sustained throughout the stimulation period. As expected, the addition of DAMGO reduced both forskolin- and A23187-stimulated fluorescence immediately and this inhibition was persistent. Quantification of both the peak and total responses indicate that DAMGO significantly decreased cAMP activity after forskolin or A23187 application.

**Figure 2.**
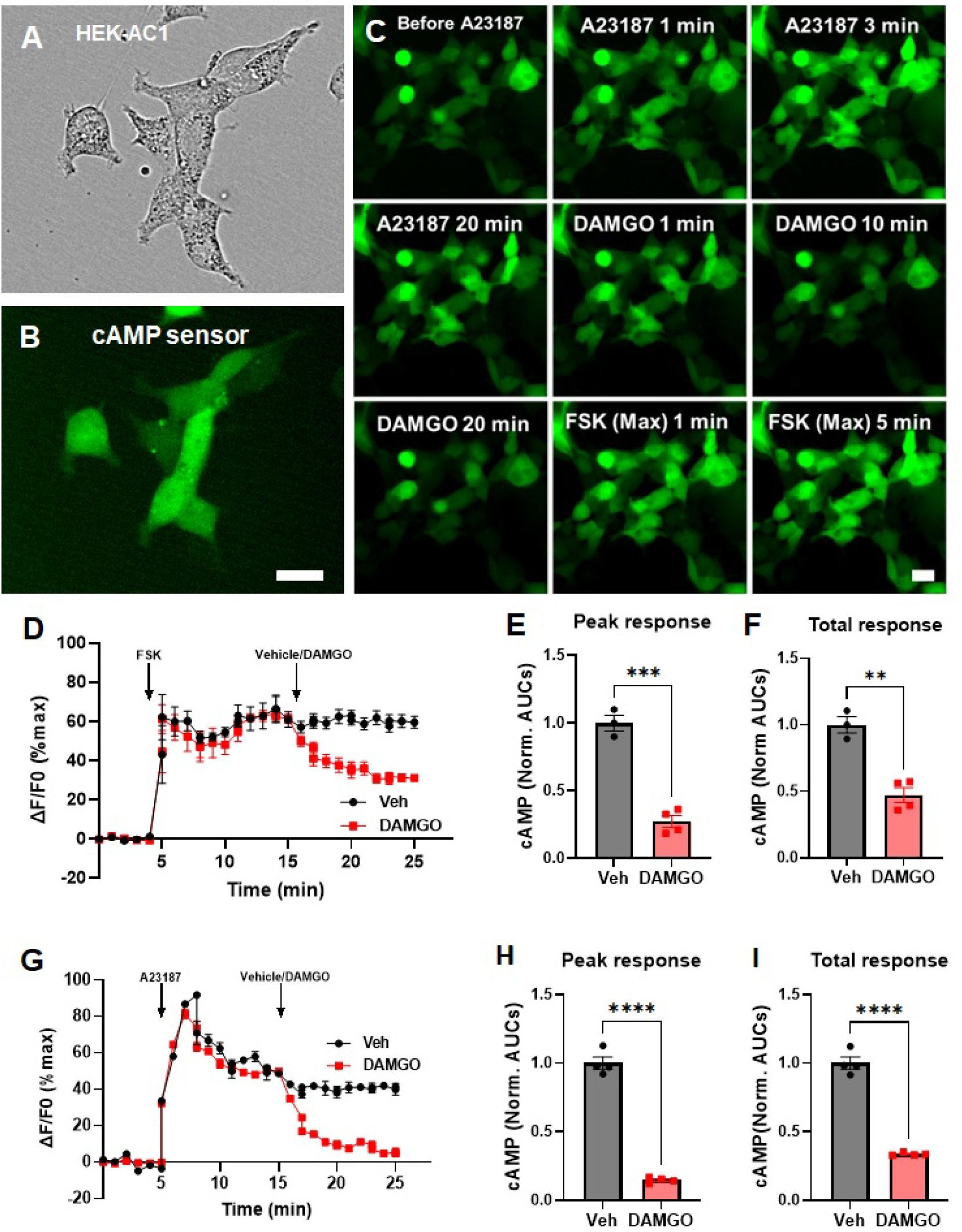
mu-opioid receptor (MOR) agonist DAMGO-mediated inhibition of cAMP activity in HEK-AC1 cells. A-B) Brightfield (A) and GFP (B) images of HEK-AC1 cells expressing MOR and cADDis cAMP biosensor. C) Time-lapse images before (pre) and after stimulation of AC1 by 1 µM A23187, subsequent inhibition of AC activity by 1 µM DAMGO and maximal (Max) AC stimulation (10 µM FSK in the presence of 100 µM IBMX and 10 µM naloxone). Scale bar: 20 µm (A-C). D-I) cAMP kinetics following stimulation with 1 µM forskolin (FSK) or 5 µM A23187 (A23) followed by the addition of DAMGO as indicated. Bar graphs showing DAMGO-mediated reduction of cAMP activity for both peak and total responses. For FSK (panel D-F) N = 3; n = 3 (Veh) and n = 4 (DAMGO), and for A23187 (panel G-I) N = 3; n = 4 (Veh) and n = 4 (DAMGO). Shapiro-Wilk test of normality, and two-tailed unpaired t-test. Statistical significance is indicated as follows: *p < 0.05, **p < 0.01, ***p < 0.001, ****p<0.0001.

### 3.3 Heterologous sensitization in HEKΔAC3/6 KO cells expressing exogenous AC1 and MOR

HEK-AC1 cells expressing the cAMP sensor and MOR were used to quantify cAMP overshoot in the absence or presence of a series pharmacological treatments. These interventions include the non-selective opioid receptor antagonist, naloxone (NAL); the Gαi/o “inhibitor”, pertussis toxin (PTX); and the neddylation inhibitor, MLN4924[34]. Initially, using the cAMP sensor, we characterized the kinetics of AC activity in response to stimulation with forskolin (FSK) after 18-hr treatment of DAMGO or vehicle (Figure 3A). Cells treated chronically with DAMGO had significantly greater cAMP activity compared to vehicle control cells (Figure 3B,C). After establishing the ability to quantify AC sensitization in HEK-AC1 cells, we used A23187 to selectively investigate heterologous sensitization of AC1 induced by chronic DAMGO treatment. Additionally, the effects of treatments with DAMGO alone, or in combination with naloxone (NAL), pertussis toxin (PTX), and MLN4924 on A23187-stimulated cAMP overshoot were evaluated. The results showed that DAMGO induced significant A23187- stimulated cAMP overshoot compared to the vehicle control, with A23187 producing a peak response within 10 minutes that stabilized within 30 minutes (Figure 3D). PTX and NAL exhibited similar efficacy in reducing A23187-stimulated cAMP overshoot, compared to DAMGO alone, while MLN4924 demonstrated a stronger inhibition, reducing the peak response by over 80% (Figure 3D-F).

**Figure 3.**
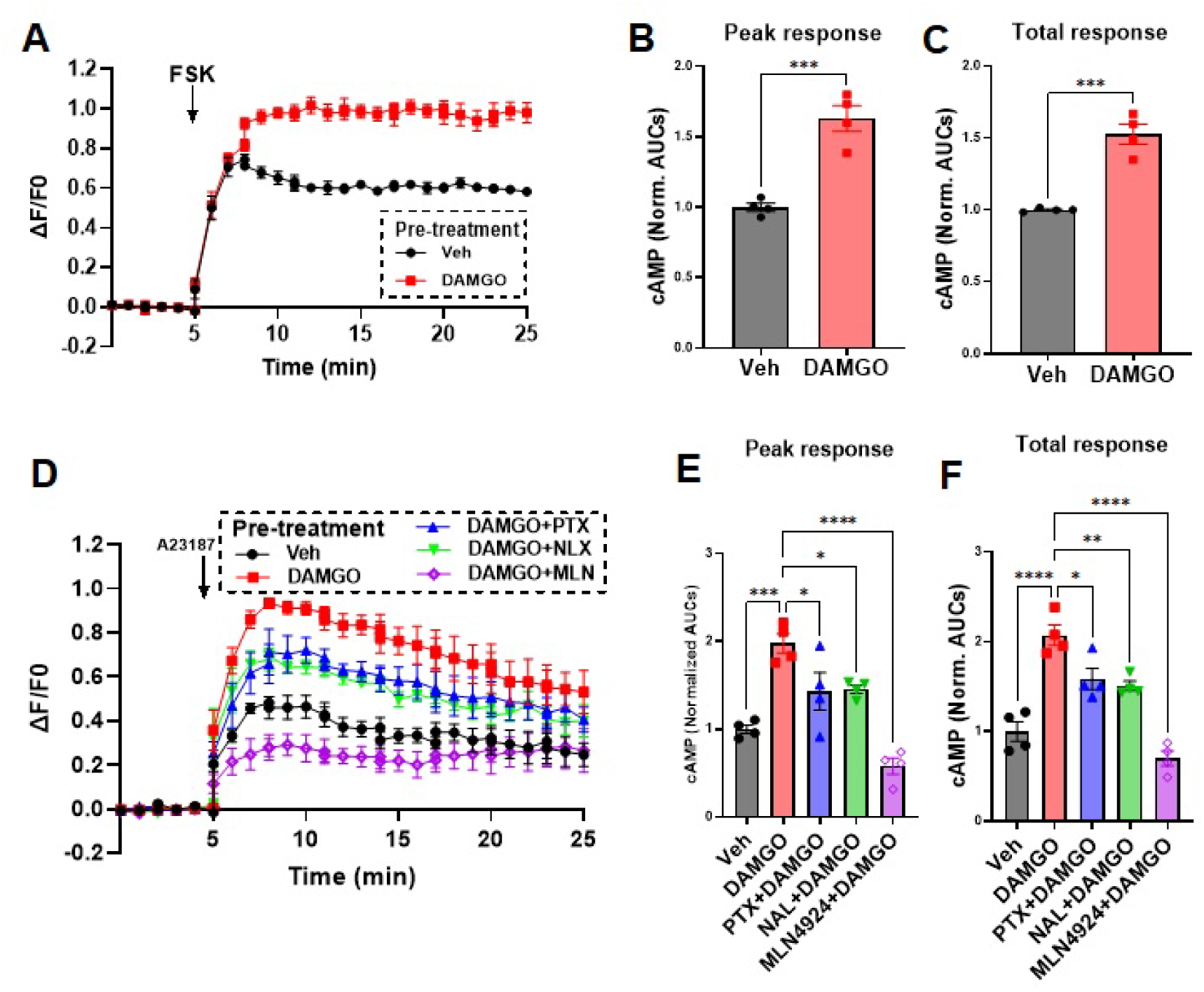
cAMP overshoot after chronic treatment with DAMGO in HEK-AC1 cells. A-C) Cells expressing cAMP-sensor and the MOR were treated with DAMGO or vehicle for 18 hr and AC activity was stimulated with 0.5 μM forskolin (FSK). Bar graphs showing cAMP overshoot peak and total responses. D-F) Cells were pretreated with vehicle or DAMGO for 18 hrs in the absence or presence of naloxone (NAL) or pertussis toxin (PTX) or MLN4924 as indicated followed by AC1 activation using A23187. Bar graphs showing peak and total cAMP overshoot responses. All error bars denote the SEM. For FSK stimulation (panel A-C), N=3; n=4 (Veh), n=4 (DAMGO); For A23187 stimulation (panel D-F) N=3; n=4 (all treatment groups). Shapiro-Wilk test of normality, followed by two-tailed unpaired t-test (panel B-C), or one-way ANOVA along with Tukey’s multiple comparisons test (panel E-F). Statistical significance is indicated as follows: *p < 0.05, **p < 0.01, ***p < 0.001, ****p<0.0001.

### 3.4 Endogenous adenylyl cyclase and MOR activity in SH-SY5Y neuroblastoma cells

SH-SY5Y neuroblastoma cells are known to express multiple adenylyl cyclase (AC) isoforms; however, the activity and kinetics of endogenous AC1 have not been well-characterized [35]. To explore this, we expressed the EPAC2-GFP cAMP-sensor in SH-SY5Y cells and stimulated cAMP activity, with either 0.5 µM forskolin or 1 µM A23187, followed by a maximal stimulation with 10 µM forskolin and 100 µM IBMX, similar to studies with the HEK-AC1 cells. Figure 4A-F shows representative time-lapse captures before and after the addition of A23187, followed by maximal stimulation using forskolin and IBMX. A significant increase in cAMP kinetics is observed upon the addition of forskolin or A23187 to SH-SY5Y cells (Figure 4G-I). The initial stimulation with either forskolin or A23187 reached a plateau within 10 minutes, with the signal sustained throughout the stimulation period. The response to A23187 was significant consistent with a Ca^2+^/CaM-stimulated AC1 response. The A23187 response is reduced compared to forskolin presumably due to the expression of other AC isoforms and differential mechanism of AC activation for Ca^2+^/CaM vs forskolin.

**Figure 4.**
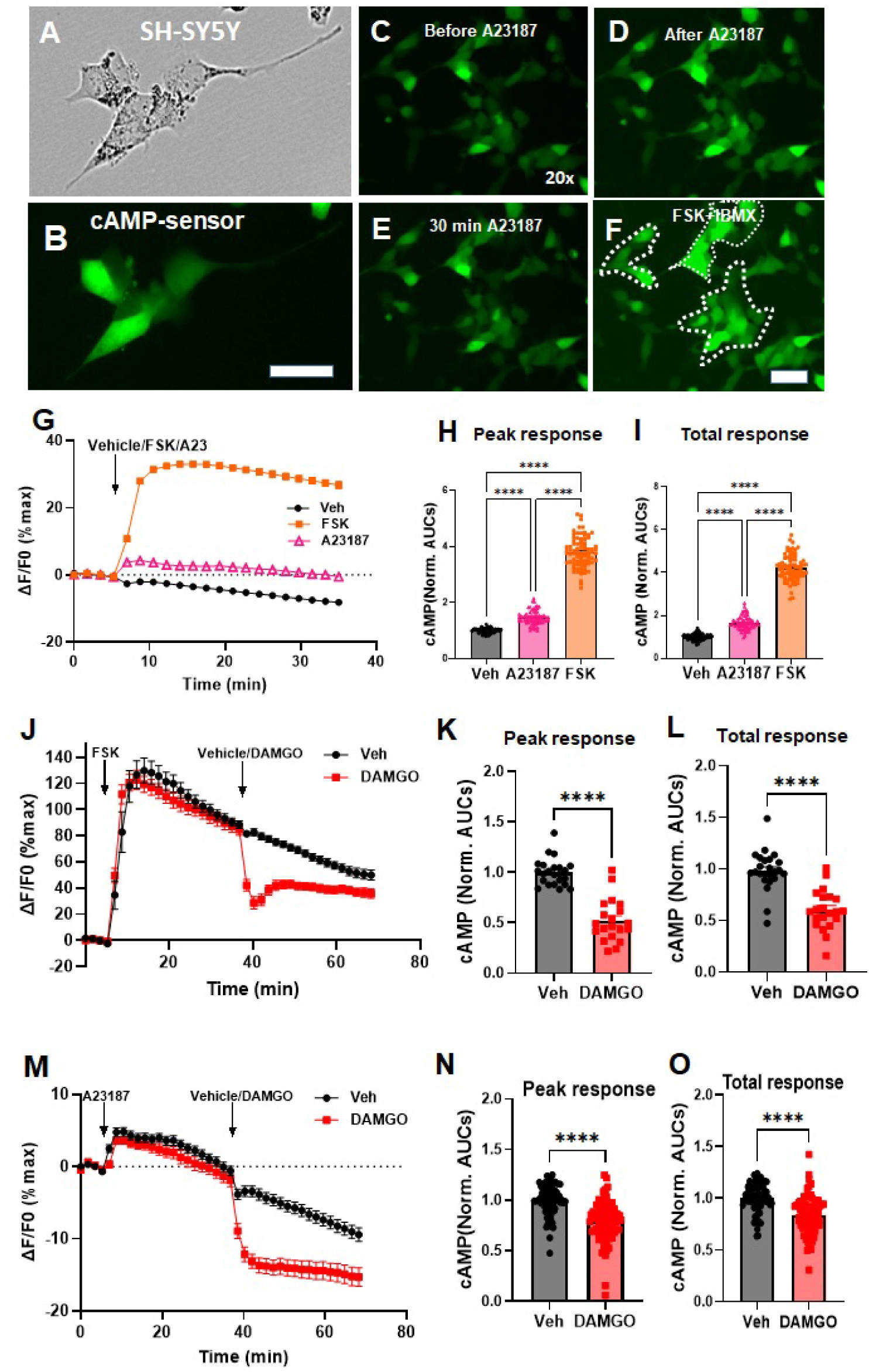
Regulation of endogenous adenylyl cyclase activity in SH-SY5Y neuroblastoma cells. A-B) Expression of cADDis cAMP biosensor in SH-SY5Y cells. C-F) Time lapse images of SH-SY5Y cells before (C) and after activation of endogenous adenylyl cyclase by 5 µM A23187 (D-E), and 10 µM FSK in the presence of IBMX (100 µM) (F). Dotted lines in panel F) indicate examples of regions of interest (ROIs) used for kinetic analysis. Scale bar: 20 µm. G) cAMP kinetics following stimulation with 5 µM A23187 (A23), 1 µM forskolin (FSK), or vehicle. Data are normalized to the maximum stimulus (FSK+IBMX). H-I) Peak and total cAMP responses after drug addition as indicated for each region of interest (ROI). J-O) Inhibition of A23187- and forskolin-stimulated cAMP by DAMGO in SH-SY5Y cells transduced with cADDis cAMP biosensor. K, L and N, O) Bar graph showing reduction of the peak and total cAMP activity by DAMGO (1 µM). All bars denote average, and all error bars denote the SEM. For panel G-I) N=3, n=86 (Veh), n=71 (A23187), n=83 (FSK), for panel J-L) N=3, n=20 (Veh), n=23 (DAMGO), and for panel M-O) N= 3, n=81 (veh), n=88 (DAMGO). Shapiro-Wilk test of Normality, followed by ordinary one way ANOVA with Tukey’s multiple comparisons test (H-I), two tailed unpaired t-test K-L), or Mann Whitney test (N-O). Statistical significance is indicated as follows: *p < 0.05, **p < 0.01, ***p < 0.001, ****p<0.0001.

To confirm the presence of endogenous MOR activity, we assessed the inhibitory effects of DAMGO on forskolin- and A23187-stimulated cAMP accumulation. We found that acute treatment with DAMGO following forskolin or A23187 stimulation resulted in a significant reduction in cAMP levels (Figure 4J-O). Specifically, after forskolin stimulation, DAMGO led to a 40% reduction in cumulative cAMP activity and a 48% reduction in peak response (Figure 4K,L). DAMGO inhibition following A23187 stimulation caused a rapid decrease in cAMP activity by ∼20% (Figure 4N,O).

### 3.5 Inhibition of endogenous and exogenous AC1 heterologous sensitization in SH-SY5Y cells

To investigate heterologous sensitization induced by chronic MOR agonist treatment by endogenously expressed AC and MOR, we again utilized SH-SY5Y cells expressing the EPAC2-GFP cAMP sensor. The aim was to assess AC1-mediated cAMP overshoot following activation with A23187. Cells were treated with DAMGO alone or following pretreatment with naloxone, PTX, or MLN4924 as described above for the HEK-AC1 cells. After drug treatment, AC1 activity was stimulated A23187 revealing a significant DAMGO-mediated cAMP overshoot compared to the control-treated cells. A23187 produced a peak response within 5 minutes that was attenuated within 20 minutes (Figure 5A). The peak and cumulative responses for A23187-stimulated cAMP overshoot following chronic DAMGO treatment were significantly elevated to 1.2-fold compared to the control. In terms of inhibition of heterologous sensitization, PTX (Gαi/o “inhibitor”) and NAL (opioid receptor antagonist) exhibited similar efficacy in reducing A23187-stimulated cAMP overshoot compared to DAMGO alone. The neddylation blocker, MLN4924 appeared to demonstrate a stronger inhibition, essentially eliminating the peak response. Because heterologous sensitization requires Gβγ subunits, we also investigated the effect of co-expressing βARK-CT, the Gβγ sequestering reagent. Transduction with βARK-CT also inhibited DAMGO-mediated cAMP overshoot (Fig. 5A-C). The modest, but significant responses to the endogenous sensitization players (i.e. MOR and AC1) prompted a similar set of experiments where both AC1 and μ-OR were overexpressed in the SH-SY5Y model background. Cells were transduced with the cAMP-sensor along with viral constructs for AC1 and MOR for sensitization experiments. Similar to the endogenous AC1 and MOR condition, DAMGO treatment sensitized A23187-stimulated cAMP in this recombinant model and the Ca^2+^/CaM-stimulated cAMP overshoot was prevented in the presence of naloxone (NAL), pertussis toxin (PTX), MLN4924, and βARK-CT (Fig. 5D-F).

**Figure 5.**
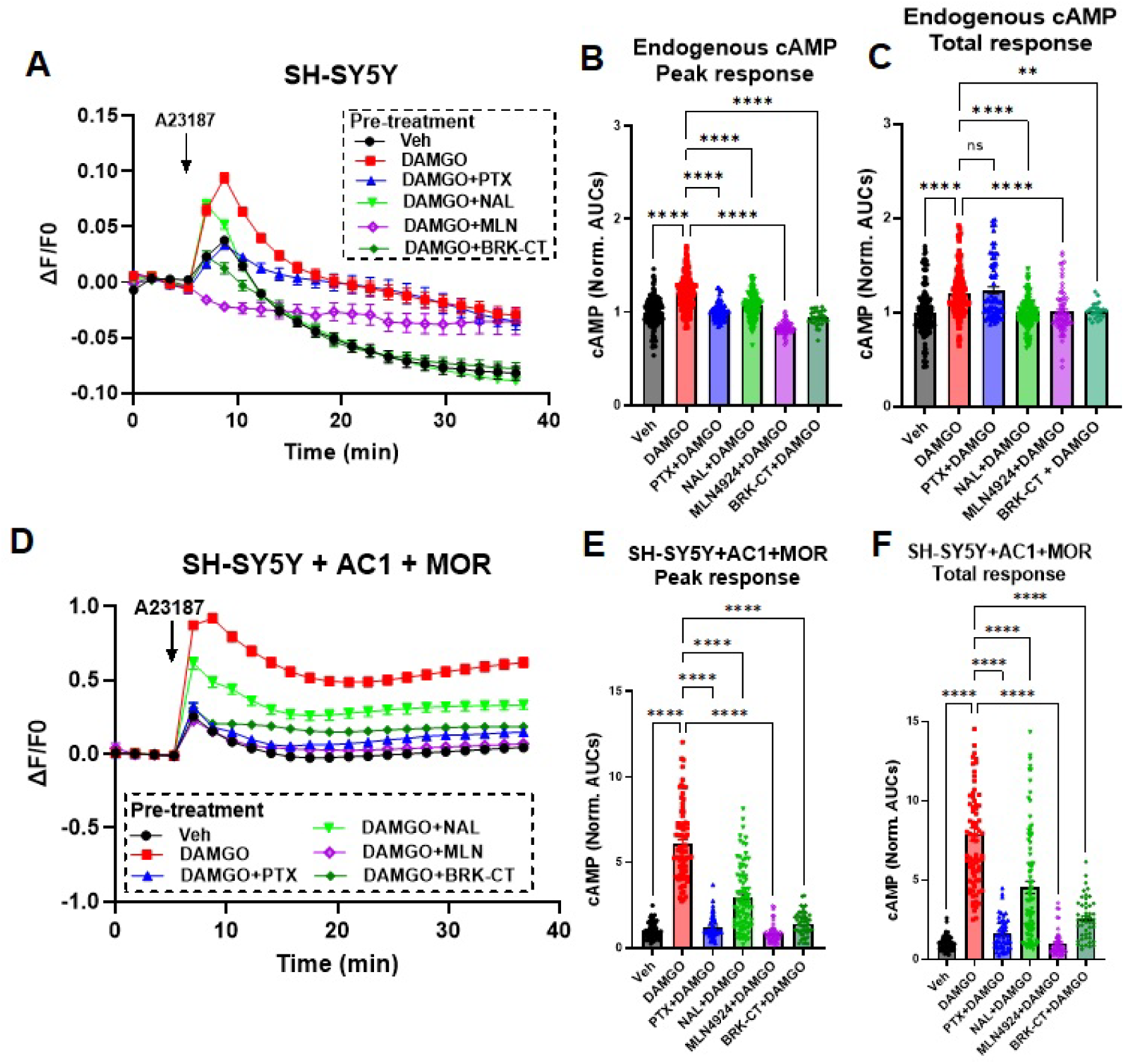
cAMP overshoot in SH-SY5Y cells stimulated with the calcium ionophore A23187. A-C) Wild-type SH-SY5Y cells or D-F) SH-SY5Y cells expressing recombinant AC1 and MOR stimulated were pretreated with vehicle or DAMGO alone for 18 hr in the absence or presence of naloxone (NAL), pertussis toxin (PTX), MLN4924 or βark-CT transduction as indicated. AC activity was stimulated using A23187 in the presence of naloxone (NAL). (A, D) Average ΔF/F traces following A23187 stimulation for wild-type (A) or recombinant AC1-expressing (D) SH- SY5Y cells. B-C and E-F) Peak and total cAMP (AUCs) for all ROIs for the A23187 response (30 minutes). All bars denote average, and all error bars denote the SEM. For panel A-C N=4, n=70-160 ROIs, and for D-F N=4, n=50-85 ROIs. Shapiro-Wilk test for normality, followed by the Kruskal-Wallis test and Dunn’s multiple comparisons test. Statistical significance is indicated as follows: *p < 0.05, **p < 0.01, ***p < 0.001, ****p<0.0001.

### 3.6 Chronic morphine exposure in SH-SY5Y cells decreases potassium flux, which is mitigated by inhibition of AC1 and EPAC

After establishing cAMP activity could be increased upon chronic stimulation with opioids *in vitro* using the EPAC2-GFP cAMP-sensor, we wanted to determine if inhibiting AC or EPAC activity would influence K_ATP_ channel activity. A thallium (Tl⁺) flux assay was used to measure the activity of potassium ions before and after exposure to the K_ATP_ channel opener diazoxide (Figure 6A-D). SH-SY5Y cells exposed to diazoxide have increased potassium flux, which is reversed by pre-exposure to glyburide, a pan-K_ATP_ channel antagonist (Figure 6E, F). Exposure to 0.5 uM morphine for 72hrs did not affect sensitivity to diazoxide, but pre-exposure to 10 uM morphine attenuated potassium flux (Figure 6E). Data presented as area under the curve illustrate a similar point compared to percent change from baseline (Figure 6F). Pre-exposure to either the EPAC1/2 inhibitor, ESI-09, or the AC1 inhibitor ST034307 for one hour, significantly elevated diazoxide-induced potassium flux in cells treated with 10 uM morphine for 72 hrs (Figure 6G). Similar results were also obtained after chronic exposure to DAMGO for 72 hrs, as ST034307 was able to reverse the inhibition of diazoxide-evoked potassium channel activity (Figure 6H).

**Figure 6.**
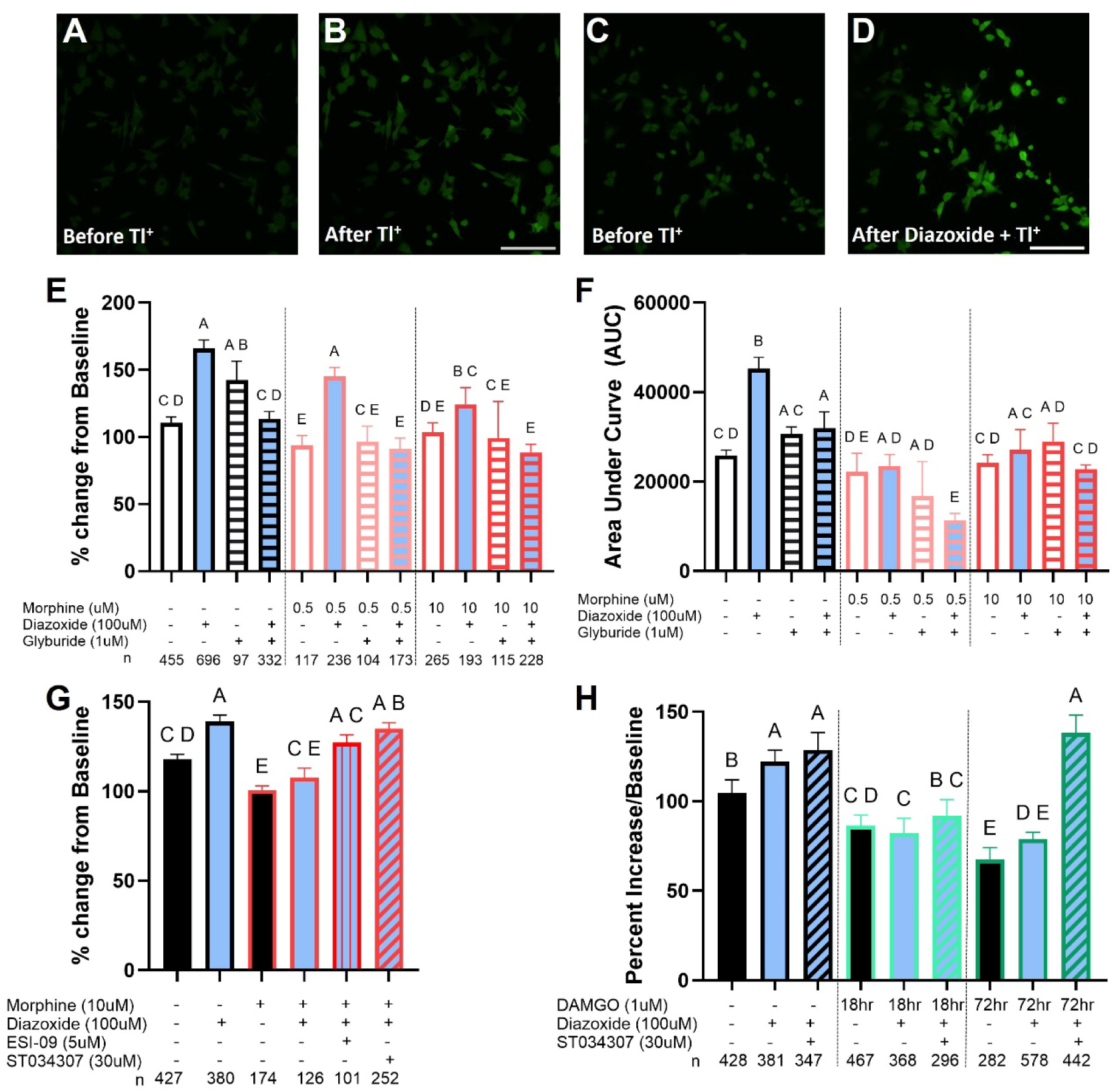
Decreased diazoxide-induced potassium flux in SH-SY5Y cells after chronic opioid stimulation. Photomicrographs of cells (A) Before thallium (Tl+) (B) After thallium (Tl+) in non-drug exposed cells, (C) Before and (D) after thallium addition in the prescence of the KATP channel opener, diazoxide (100uM). (E) Potassium flux measurements comparing the peak response compared to baseline, using thallium-based assay and confocal microscopy on SH-SY5Y cells. Exposure to diazoxide, a SUR1-selective KATP channel subtype agonist, increases potassium flux compared to baseline. Prior exposure of a KATP channel antagonist, glyburide, decreases potassium flux back untreated levels. Chronic administration of morphine at 0.5 uM and 10 uM over 72 hours, also decreases potassium flux. (F) Data in A presented as area under the curve. (G) Pre-exposure to a EPAC inhibitor, ESI-09, significantly increases potassium flux after morphine treatment (10uM, 72 hrs). (H) Inhibition of AC1 by pre-exposure to ST034307 significantly increases potassium flux compared to diazoxide treatment alone after chronic morphine exposure for 72 hrs and (D) chronic DAMGO exposure for 18 or 72 hrs. Data presented as median ± 95% CI from individual cells from three separate imaging experiments. Images in A-D obtained at 40x using a Nikon spinning disc confocal microscope, Scale bar = 100uM. Significance determined using Kruskal-Wallis test with Dunn’s post-hoc test, groups lacking a common letter are significantly different (P<0.05).

### 3.7 K_ATP_-induced potassium flux is decreased in mouse DRG after chronic opioid exposure, which is largely reversed by inhibition of EPAC2

Potassium channel activity was also measured in primary DRG cultures from mice chronically treated with morphine (15mg/kg, 5 days, twice daily) or controls. DRG cultures were plated overnight and loaded with thallium-indicating dye similar to immortalized cell cultures (Figure 7A,B), but were also co-labeled for the presence of ChBTx and/or IB4 (Figure 7C). Overall, DRGs were sensitive to diazoxide, but this potassium flux was eliminated after pre-incubation with glyburide (Figure 1D). Incubation with HJC0350, a partially selective inhibitor for EPAC2 compared to EPAC1, was able to largely reverse the inhibition of diazoxide-induced potassium channel activity after chronic morphine exposure (Figure 7D). HJC0350 also increased potassium channel activity in myelinated cells (ChBTx^+^ cells, Figure 7E), IB4+ cells (Figure 7F), and DRG without labels (Figure 7G). HJC0350 and ST034307 both significantly elevated potassium flux in morphine-treated mice in DRG positive for both ChBTx and IB4 (Figure 7H).

**Figure 7:**
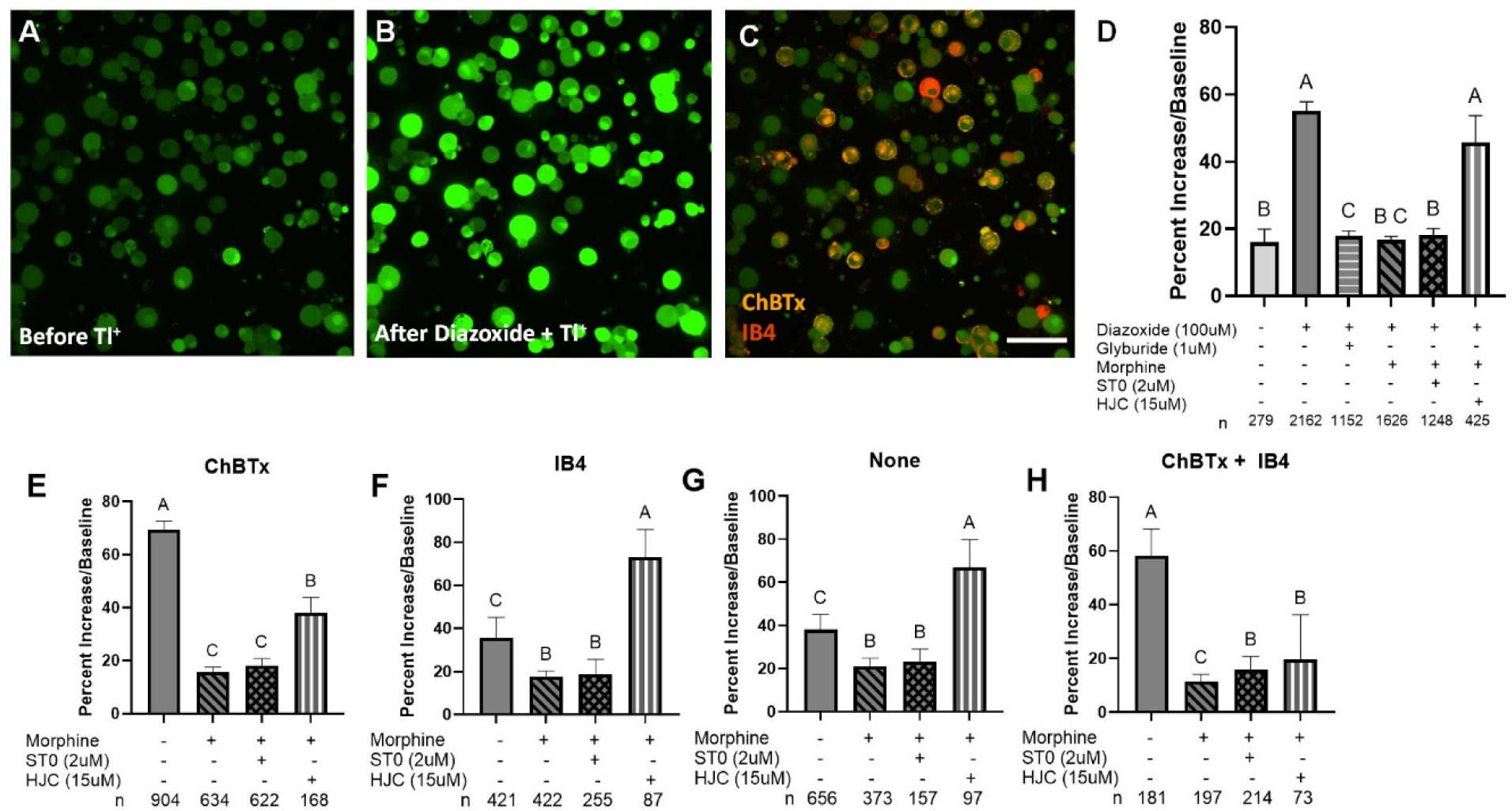
Blocking EPAC activity increases potassium flux indorsal root ganglia from naïve and morphine treated mice. (A) Thallium indicating dye is loaded into mouse DRG upon exposure to thallium will increase in intensity as potassium channels open. (B) Exposure to thallium, a surrogate for potassium ions, increases fluorescence intensity which is further enhanced by co-administration of potassium channel agonists (Diazoxide, 100uM) (C) DRG co-labeled using commercially available markers for isolectin B4 (IB4,#132450, Invitrogen) or Cholera B Toxin (ChBTx, #C22843, Invitrogen). (D) Diazoxide added to thallium significantly enhanced the peak fluorescence which is blocked by pre-treatment with glyburide (1uM) a KATP channel antagonist. Morphine treatment for 72 hrs (10uM) significantly decreases potassium flux, which is mostly reversed after pre-treatment with an EPAC antagonist, HJC 0350 (HJC). (E-H) Separation of the data from D by co-labeling for myelinated fibers using ChBTx (E), GDNF-dependent non-peptidergic C-fibers (F), and DRG that label for both markers (G). (H) DRG that do not label for IB4 or ChBTx are presumably peptidergic C-fibers. HJC 0350 significantly enhanced potassium flux in all four groups of DRG, while ST03407 significantly increased potassium flux in DRG co-labeled for ChBTx and IB4 (G). Data presented as change in fluorescence intensity compared to baseline before thallium addition as medians ± 95% CI from at least three separate imaging experiments. Images obtained at 40x using a Nikon TiS inverted microscope. Scalebar = 100uM. Kruskal-Wallis test with Dunn’s post-hoc test, groups lacking a common letter are significantly different (P<0.05).

## 4. Discussion

Current treatments for chronic pain predominantly rely on opioids, which offer substantial efficacy in pain relief but are associated with significant risks, including the development of pharmacological tolerance, physiological dependence, and addiction [36, 37]. Chronic opioid exposure also leads to an increase in cAMP activity, presumably through hyperactivation of adenylyl cyclases in the peripheral and central nervous system. ACs mediates the production of cyclic AMP (cAMP), a key secondary messenger involved in various physiological processes, including pain perception [38]. In this study, we have identified used both *in vitro* and *in vivo* approaches to implicate EPAC2 as a potential molecular regulator downstream of AC1 that acts as a brake on K_ATP_ channels.

Loss of AC1 activity and expression has been found to reduce hypersensitivity in chronic pain models, and reduce opioid-induced tolerance in several rodent models [31, 39, 40]. Global knockout of AC1 reduces formalin-induced nocifensive responses and mechanical allodynia after Complete Freund’s Adjuvant injection [41]. The activity of adenylyl cyclases contributes to hypersensitivity in the brain, but also in the spinal cord, as intrathecal injections of forskolin produces mechanical allodynia in both hindpaws, but not in AC1 and AC8 double knock out (DKO) mice [42]. AC1 KO mice treated chronically with morphine using an escalating dose paradigm, have significantly lower withdrawal scores compared to wildtype littermates [43]. These previous reports are consistent with the data collected here using a viral strategy to knockdown AC1 in the spinal cord and DRG, where we observed decreased morphine tolerance and withdrawal. The utility of therapeutics targeting adenylyl cyclases, particularly AC1 may prove to have great utility in the future [39]. For example, when given systemically, ST034307 was able to rescue acid-depressed mouse nesting behaviors and abdominal constrictions in a lactic acid model of visceral pain, which may be due to its ability to reduce cAMP concentration in mouse DRG [44].

This research aimed to overcome the limitations of traditional *in vitro* models of opioid signaling by developing a sensitive and reliable neuronal model using SH-SY5Y neuroblastoma cells expressing genetically encoded EPAC2-GFP cAMP biosensors. These biosensors enabled real-time monitoring of cAMP dynamics in live cells, providing a powerful tool for studying the sensitization of endogenous adenylyl cyclase activity mediated by Gαi/o-coupled receptors. Consistent with a Gαi/o-coupled receptor-dependent mechanism, pretreatment with the opioid receptor antagonist naloxone (NAL) or inactivation of Gαi/o proteins with pertussis toxin (PTX) prevented DAMGO-mediated cAMP overshoot of endogenous adenylyl cyclase in SH-SY5Y cells [4]. Similarly, the overshot was prevented by expression of the Gβγ-sequestering reagent, βARK-CT as previously observed for AC1[45]. These observations extend previous studies in recombinant models to a human neuronal model and offer insights into the molecular mechanisms of cAMP sensitization during chronic opioid exposure. Ultimately, these experiments provided a robust model for investigating both endogenous and exogenous AC1 activity in neuronal cells while having the advantage of identifying effective inhibitors to mitigate cAMP overshoot in the future.

The ability of the neddylation inhibitor, MLN494 to prevent DAMGO-mediated cAMP overshoot is an important observation and consistent with our studies in recombinant HEK cell models and native NG108-15 cells [46]. The involvement of the neddylation pathway in heterologous sensitization was discovered as part of a genome-wide siRNA screen that revealed the involvement of several candidates for cAMP overshoot following chronic D_2_ dopamine receptor activation, including: cullin-3 protein (CUL3), RING-box protein 1, Neural precursor cell expressed developmentally downregulated protein 8 (NEDD8), and a NEDD8-activating enzyme. Subsequent *in vivo* studies revealed the involvement of the neddylation (via MLN4924) in locomotor sensitizing effects of ethanol and CFA-induced allodynia [47]. More recently, MLN4924 was shown to reduce opioid-induced cAMP overshoot and dependence *in vivo* [48], and our results using the EPAC2-GFP cAMP-sensor match these results *in vitro*. Treatment with MLN4924, or the proteasome inhibitor bortezomib, prevents heterologous sensitization suggesting the premise that chronic MOR stimulation involves the loss of expression or negative regulation of a key player in these pathways, including neddylation activating Cullin RING ligases (CRLs), which promote the ubiquitylation and degradation of proteins. Such a player could be involved with overactivity of AC1 and/or EPAC2 and/or the loss of potassium channel activity. Specifically with regards to K_ATP_ channels, treatment of COS cells with proteasome inhibitors leads to increased surface levels of transiently expressed SUR1 and Kir6.2 [49], and diazoxide’s efficacy to reduce somatostatin release in TGP52 cells is increased after shRNA- mediated cullin protein knockdown[50].

Many cAMP-mediated cellular effects directly or indirectly contribute to long-term changes in neuronal excitability, including synaptic transmission. Protein kinase A (PKA), another downstream target of cAMP, has also been implicated in tolerance and withdrawal. Central administration of antagonists or antisense oligodeoxynucleotides targeting of PKA completely blocked withdrawal and tolerance from morphine in rodent models [51, 52]. The relationship between PKA and K_ATP_ channels is complicated, it appears that Kir6.2/SUR1 subtypes, the predominant subtype found in the peripheral and central nervous system, may be either inhibited in the presence of PKA [53] or activated [54] depending on additional factors such as ATP/ADP levels. We previously did not find any upregulation of the catalytic or regulatory subunits of PKA in mouse spinal cord or DRG after chronic morphine administration, but Rapgef4 (protein: EPAC2) was significantly increased [31]. Although other isoforms of ACs have been implicated in morphine tolerance [55], our data are in agreement with previous studies that AC1 is a contributing factor in the development of opioid tolerance and withdrawal. Our study does not rule out the possibility of alternative AC isoforms involved in inhibition of potassium channels or other ion channels contributing to opioid signaling and hypersensitivity in the nervous system.

After establishing heterologous sensitization using *in vitro* models, we were able to more closely examine the effects of AC or EPAC inhibition on potassium channel activity. The AC1 inhibitor ST034307, or the EPAC inhibitor ESI-09 both restored potassium channel activity in SH-SY5Y cells chronically treated with opioids. A similar phenomenon was observed in mouse DRGs after chronic treatment with morphine for five days. In our hands, the EPAC2 selective inhibitor HJC0350 was able to completely reverse the inhibition of diazoxide-evoked potassium activity in DRG from morphine tolerant mice. Several DRG subtypes were affected by chronic morphine exposure, as the diazoxide-induced reduction in potassium channel activity was significantly reduced in peptidergic C-nociceptors, non-peptidergic C-nociceptors, and myelinated cells. The ability for HJC0350 to reverse the reduction in potassium flux was most evident in IB4+ cells and cells without labels for either IB4 or ChBtx (presumably peptidergic C-nociceptors). The latter finding is corroborated by studies in humans, indicating that peptidergic C-nociceptors are positive for OPRM1, the MOR gene[56]. Combined, these data suggest that loss of potassium channel activity after long-term opioid exposure [57] can be reversed by inhibiting cAMP activity.

Previously, work from our laboratory and others have implicated K_ATP_ channels in opioid-induced antinociception, which decreases after chronic opioid exposure [58]. Our strategy to reduce AC1 and upregulate K_ATP_ channel subunits in the spinal cord worked to cooperatively reduce morphine tolerance and withdrawal in mice. cAMP has been implicated in modulating potassium channel currents in several previous studies [59], including inhibition of K_ATP_ channels in neuronal and non-neuronal systems [53, 60–62]. Elevation of cAMP, and thereby EPAC2 protein activity, could be an important negative regulator of potassium channels during chronic opioid exposure. EPAC2 helps to promote insulin secretion in pancreatic beta cells through Kir6.2 and SUR1 subtype K_ATP_ channels either directly through engagement through SUR1-subunits or by recruitment of Rap1. In addition, mice lacking EPAC2 have impaired ability for SUR1-targeting antagonists to close K_ATP_ channels [63] and genetic deletion of EPAC proteins increases the open probability of K_ATP_ channels [64]. The balance between PKA and EPAC signaling pathways, may predominate under the acute versus chronic pain states, respectively, and the underlying mechanisms how the EPAC pathway enhances pain signals and produces hypersensitivity is worth further consideration[65].

In conclusion, this work demonstrates that inhibition of adenylyl cyclase 1 (AC1) and EPAC can effectively restore ATP-sensitive potassium channel activity following chronic opioid exposure. This study provides evidence for targeting AC1 in the context of opioid tolerance and withdrawal, while also highlighting potential additional targets to prevent cAMP overshoot during chronic opioid exposure. Pharmacological modulation of EPAC, directly at AC1 or downstream of AC1 in the opioid receptor pathway, may overcome the limitations of previous therapeutics currently in development for opioid use disorders and/or chronic pain by providing more improved treatment options[65, 66].

## Acknowledgements

The authors would like to thank Amanda Fowler and Jacob DeHaan for their technical assistance. This work is supported by Purdue University, and NIH grant numbers R01DA051876 (AHK and VJW) and R01NS119917 (VJW).

## Supplemental Materials

**Supplemental Figure 1.**
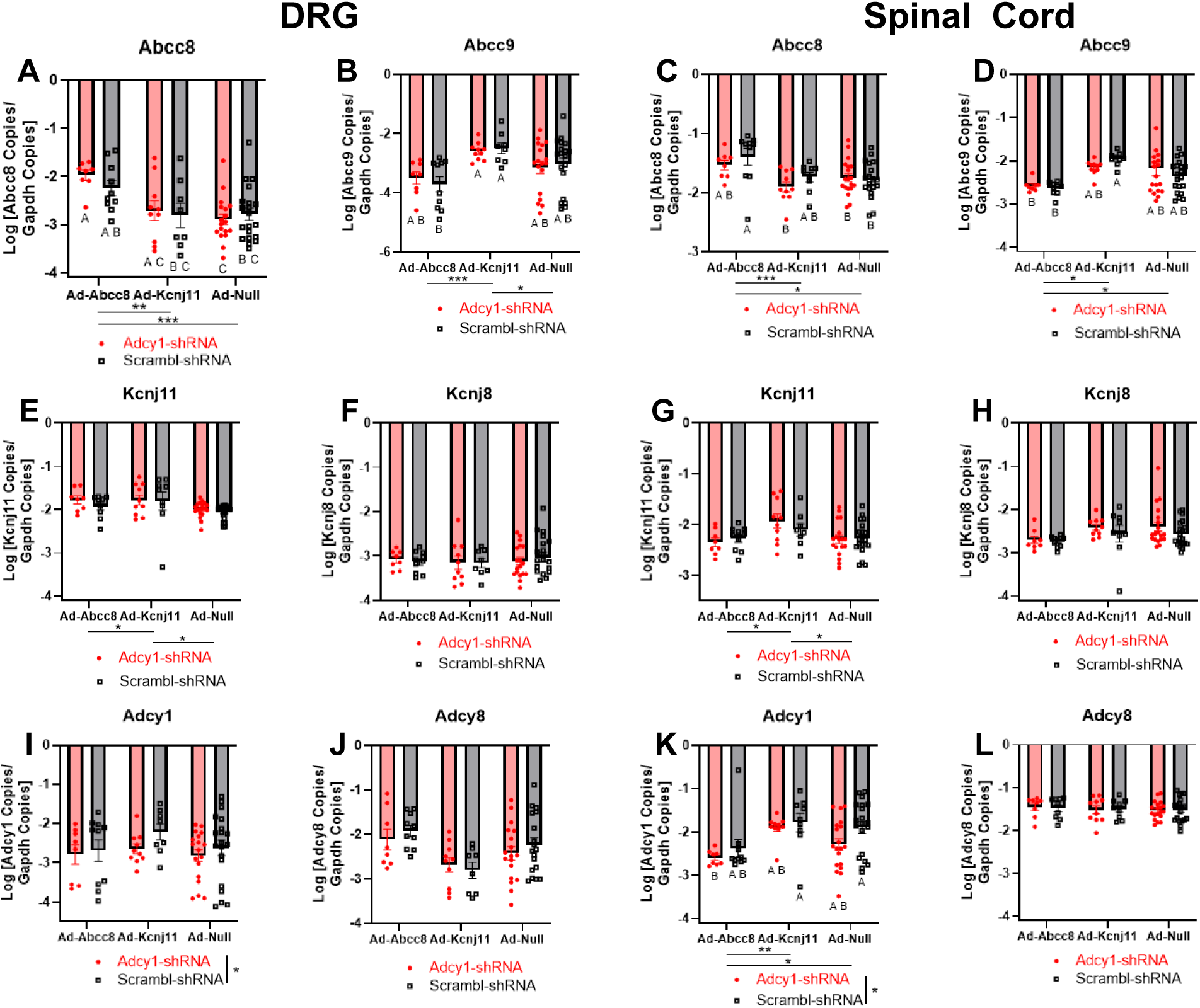
Quantification of Adcy1 knockdown and K_ATP_ channel subunit upregulation in behavioral experiments. Mice were injected with either AAV9-Adcy1-shRNA via intrathecal injection four weeks prior to behavioral experiments to downregulate AC1 (Adcy1), while other mice received AAV9-Scramble-shRNA as a control. One week before behavioral experiments, mice were injected with an adenovirus to upregulate either SUR1 (Abcc8), Kir6.2 (Kcnj11), or a viral control (Null). Gene expression of KATP channel subunits (A-H) and Adcy1 and Adcy8 were examined in the dorsal root ganglia (DRG) and spinal cord and compared across treatment groups. A. Upregulation of Abcc8 (Ad-Abcc8) significantly increased expression of Abcc8 in DRG compared to upregulation of Kcnj11 (Ad-Kcnj11) or control virus (Ad-Null). B. Abcc9 expression was significantly increased in mice with upregulation of Ad-Kcnj11 compared to Ad-Abcc8 or Ad-Null. C. Abcc8 expression was also significantly elevated in Ad-Abcc8 mice compared to Ad-Kcnj11 or Ad-Null mice. D. Similar to DRGs, Abcc9 expression was significantly increased in mice with upregulation of Ad-Kcnj11 compared to Ad-Abcc8 or Ad-Null mice. E. Kcnj11 was significantly upregulated in Ad-Kcnj11 mice compared to Ad-Abbc8 and Ad-Null mice, but F. Kcnj8 expression was not significantly altered. G. Similar expression patterns in the spinal cord were also found for Kcnj11 and H. Kcnj8 in the spinal cord. I. Adcy1 was significantly downregulated in the DRG of mice treated with Adcy1-shRNA. J. Adcy8 expression was not altered in the DRG. K. Adcy1-shRNA also significantly decreased Adcy1 expression in the spinal cord. L. Adcy8 expression was not altered in spinal cord regardless of viral treatment(s). Data were log transformed and presented as mean ± SEM. Two-way ANOVA with Tukey post-hoc test, groups lacking a common letter are significantly different (*P<0.05).

**Supplemental Figure 2.**
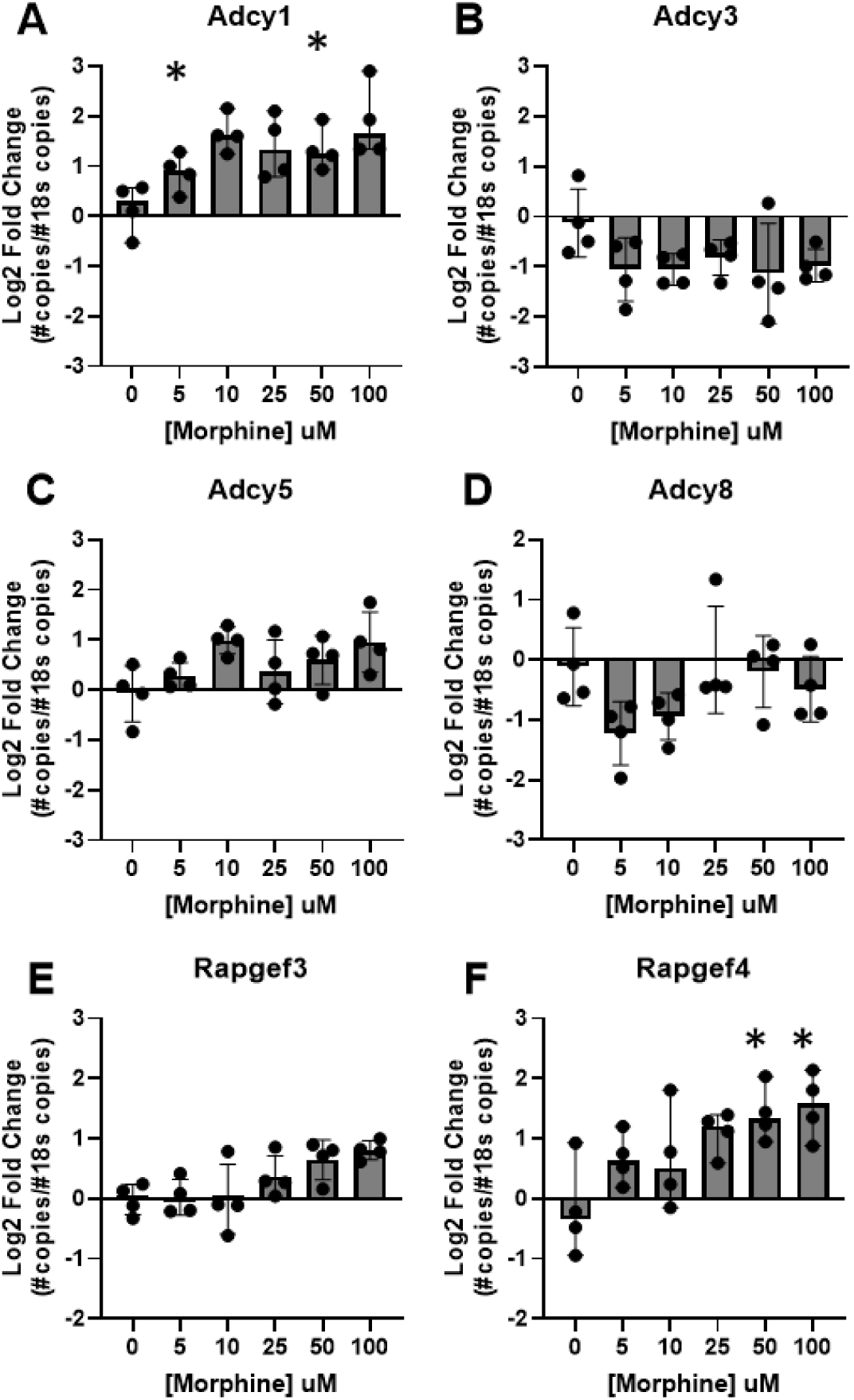
Quantification of adenylyl cyclase and Epac isoforms in SH-SY5Y cells with or without morphine for 72 hours in culture. mRNA expression of adenylyl cyclase isoforms (Adcy) and Epac1 (Rapgef3) and Epac2 (Rapgef4) quantified by qPCR for untreated cells (0 uM) versus 72 hr exposure to morphine at 5, 10, 25, 50, and 100uM. A. Adcy1, B. Adcy3, C. Adcy5, D. Adcy8, E. Rapgef3, and F. Rapgef4. Significant increases in expression were seen for Adcy1 and Rapgef4. Data presented as median ± 95% CI. Kruskal-Wallis test with Dunn’s post-hoc test compared to 0 uM morphine,*P<0.05.

**Supplemental Table 1.**
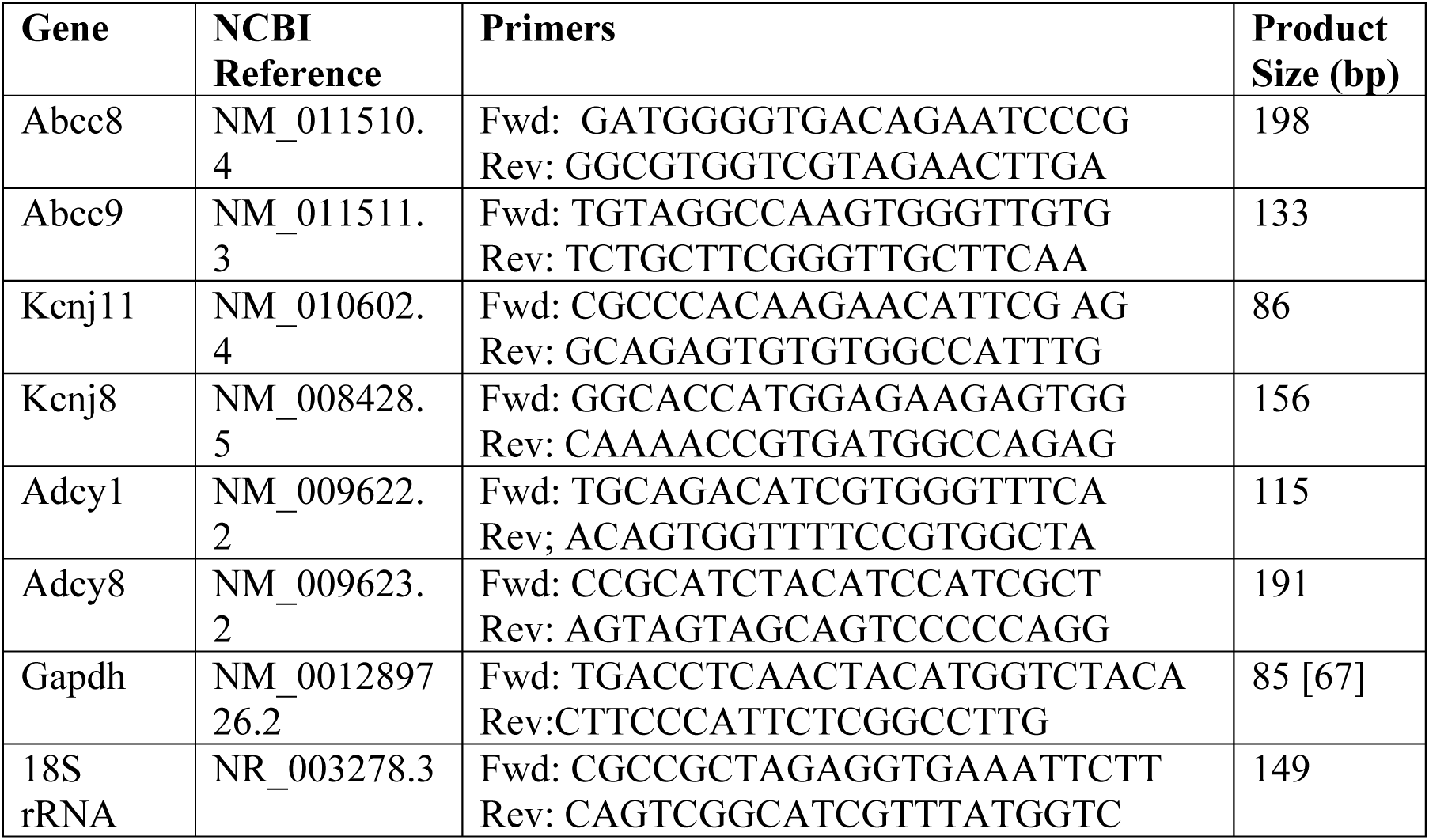
Names, sequences and amplicon lengths of primers for mouse housekeeping and target genes.

**Supplemental Table 2.**
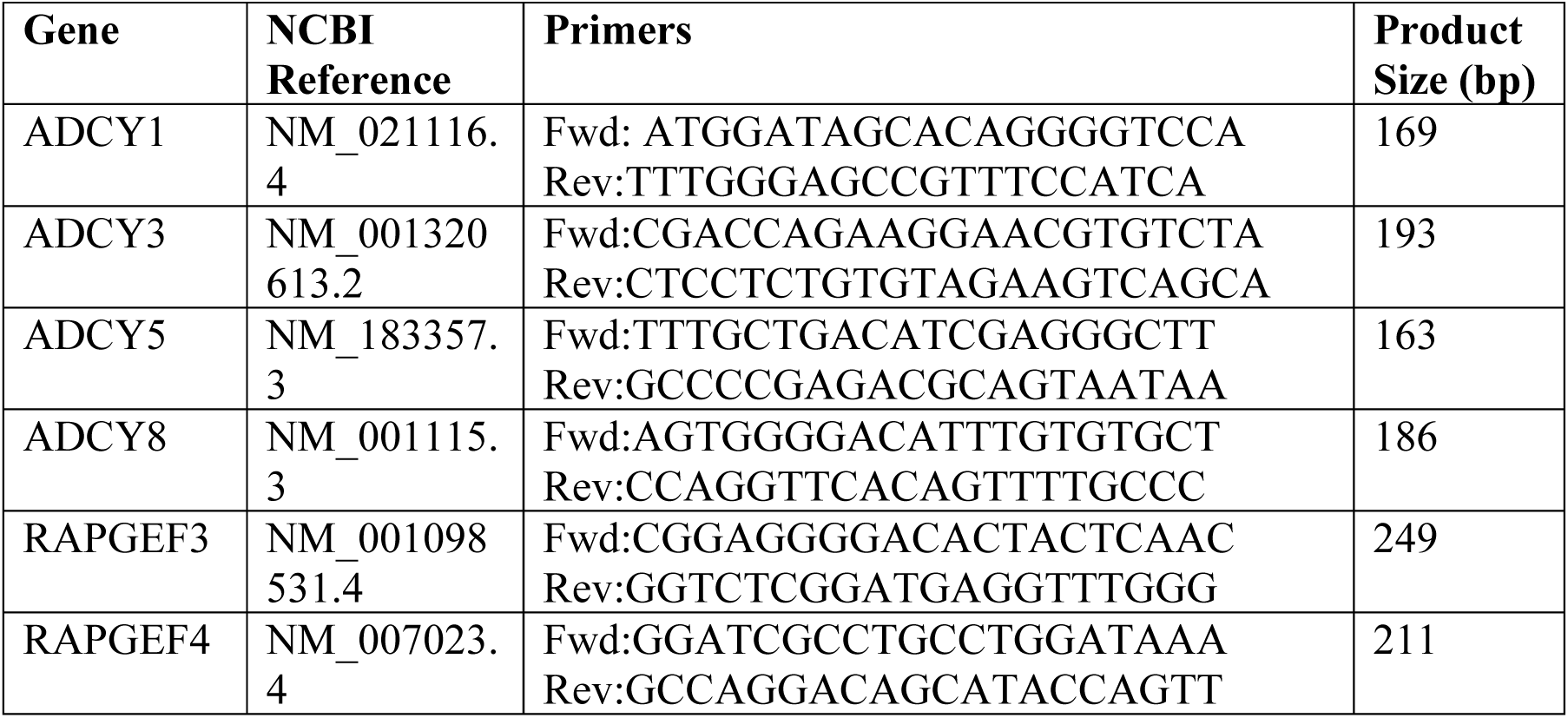
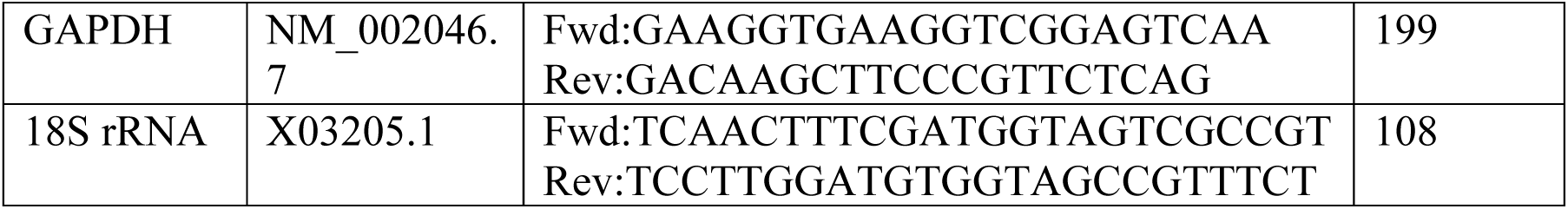
Names, sequences and amplicon lengths of primers for human housekeeping and target genes.

