## Supplemental Materials for "Inhibition of adenylyl cyclase 1 (AC1) and exchange protein directly activated by cAMP (EPAC) restores ATP-sensitive potassium (K_ATP_) channel activity after chronic opioid exposure"

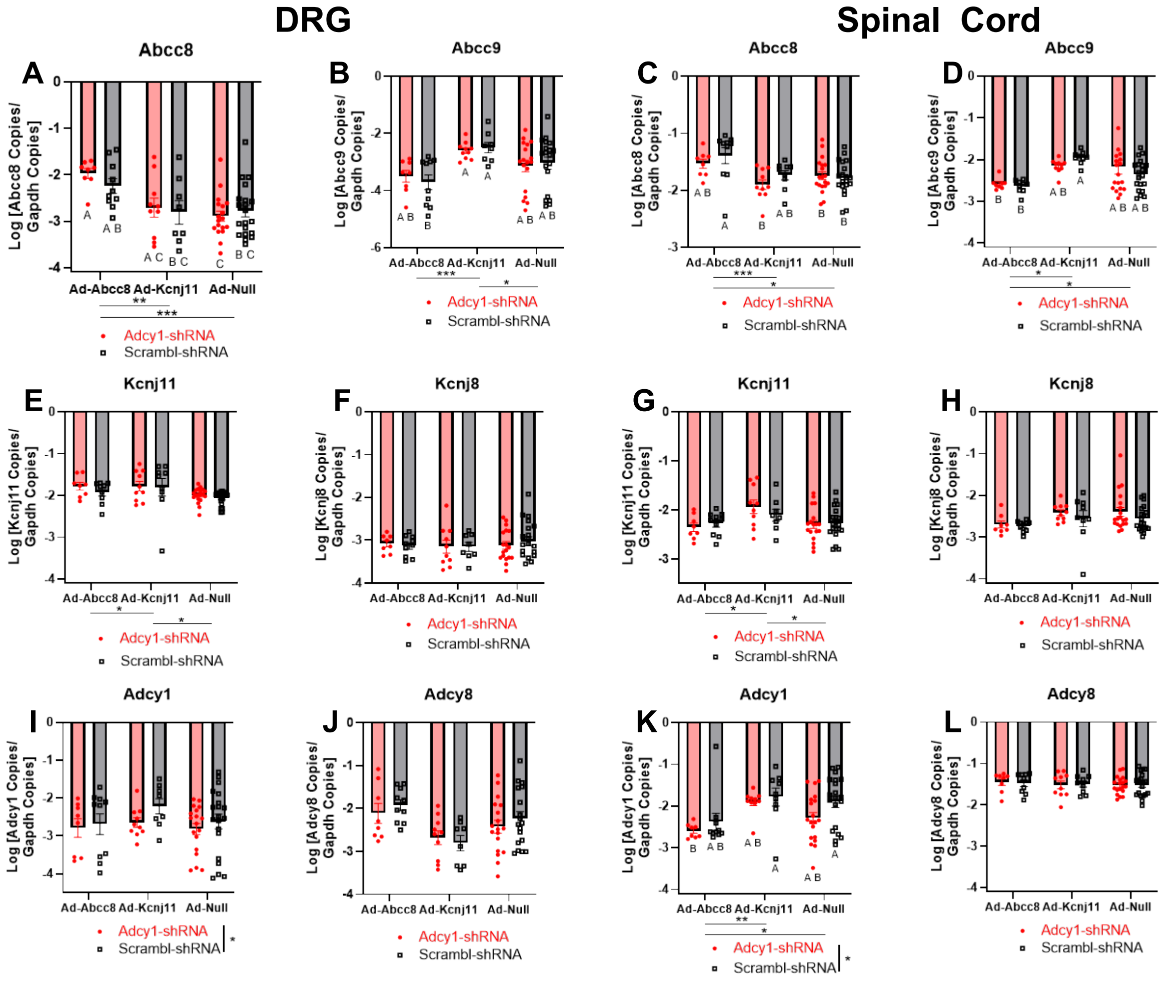


**Supplemental Figure 1. Quantification of Adcy1 knockdown and K<sub></sub>ATP channel subunit upregulation in behavioral experiments.** Mice were injected with either AAV9-Adcy1-shRNA via intrathecal injection four weeks prior to behavioral experiments to downregulate AC1 (Adcy1), while other mice received AAV9-Scramble-shRNA as a control . One week before behavioral experiments, mice were injected with an adenovirus to upregulate either SUR1 (Abcc8), Kir6.2 (Kcnj11), or a viral control (Null). Gene expression of KATP channel subunits (A-H) and Adcy1 and Adcy8 were examined in the dorsal root ganglia (DRG) and spinal cord and compared across treatment groups. A. Upregulation of Abcc8 (Ad-Abcc8) significantly increased expression of Abcc8 in DRG compared to upregulation of Kcnj11 (Ad-Kcnj11) or control virus (Ad-Null). B. Abcc9 expression was significantly increased in mice with upregulation of Ad-Kcnj11 compared to Ad-Abcc8 or Ad-Null. C. Abcc8 expression was also significantly elevated in Ad-Abcc8 mice compared to Ad-Kcnj11 or Ad-Null mice. D. Similar to DRGs, Abcc9 expression was significantly increased in mice with upregulation of Ad-Kcnj11 compared to Ad-Abcc8 or Ad-Null mice. E. Kcnj11 was significantly upregulated in Ad-Kcnj11 mice compared to Ad-Abbc8 and Ad-Null mice, but F. Kcnj8 expression was not significantly altered. G. Similar expression patterns in the spinal cord were also found for Kcnj11 and H. Kcnj8 in the spinal cord. I. Adcy1 was significantly downregulated in the DRG of mice treated with Adcy1-shRNA. J. Adcy8 expression was not altered in the DRG. K. Adcy1-shRNA also significantly decreased Adcy1 expression in the spinal cord. L. Adcy8 expression was not altered in spinal cord regardless of viral treatment(s). Data were log transformed and presented as mean ± SEM . Two-way ANOVA with Tukey post-hoc test, groups lacking a common letter are significantly different (*P<0.05).

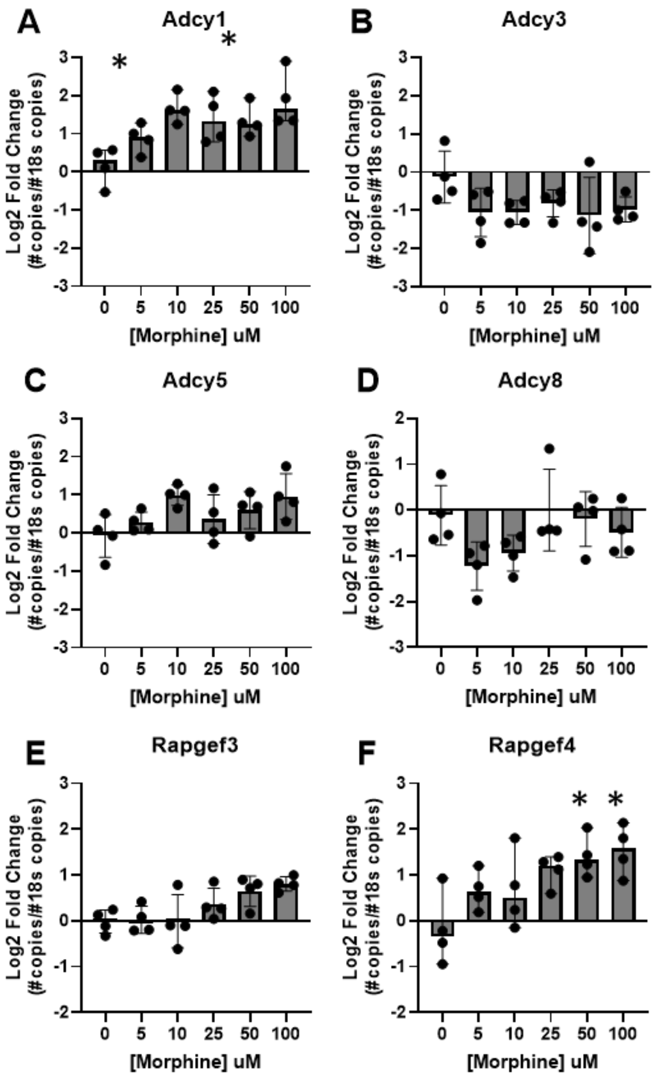


**Supplemental Figure 2. Quantification of adenylyl cyclase and Epac isoforms in SH-SY5Y cells with or without morphine for 72 hours in culture.** mRNA expression of adenylyl cyclase isoforms (Adcy) and Epac1 (Rapgef3) and Epac2 (Rapgef4) quantified by qPCR for untreated cells (0 uM) versus 72 hr exposure to morphine at 5, 10, 25, 50, and 100uM. A. Adcy1, B. Adcy3, C. Adcy5, D. Adcy8, E. Rapgef3, and F. Rapgef4. Significant increases in expression were seen for Adcy1 and Rapgef4. Data presented as median ± 95% CI. Kruskal-Wallis test with Dunn's post-hoc test compared to 0 uM morphine,*P<0.05.

**Supplemental Table 1. Names, sequences and amplicon lengths of primers for mouse housekeeping and target genes**

| Gene | NCBI Reference | Primers | Product Size (bp) |
|---|---|---|---|
| Abcc8 | NM_011510.4 | Fwd: GATGGGGTGACAGAATCCCG<br>Rev: GGCGTGGTCGTAGAACTTGA | 198 |
| Abcc9 | NM_011511.3 | Fwd: TGTAGGCCAAGTGGGTTGTG<br>Rev: TCTGCTTCGGGTTGCTTCAA | 133 |
| Kcnj11 | NM_010602.4 | Fwd: CGCCCACAAGAACATTCG AG<br>Rev: GCAGAGTGTGTGGCCATTTG | 86 |
| Kcnj8 | NM_008428.5 | Fwd: GGCACCATGGAGAAGAGTGG<br>Rev: CAAAACCGTGATGGCCAGAG | 156 |
| Adcy1 | NM_009622.2 | Fwd: TGCAGACATCGTGGGTTTCA<br>Rev; ACAGTGGTTTTCCGTGGCTA | 115 |
| Adcy8 | NM_009623.2 | Fwd: CCGCATCTACATCCATCGCT<br>Rev: AGTAGTAGCAGTCCCCCAGG | 191 |
| Gapdh | NM_001289726.2 | Fwd: TGACCTCAACTACATGGTCTACA<br>Rev:CTTCCCATTCTCGGCCTTG | 85 [67] |
| 18S rRNA | NR_003278.3 | Fwd: CGCCGCTAGAGGTGAAATTCTT<br>Rev: CAGTCGGCATCGTTTATGGTC | 149 |

**Supplemental Table 2. Names, sequences and amplicon lengths of primers for human housekeeping and target genes**

| Gene | NCBI Reference | Primers | Product Size (bp) |
|---|---|---|---|
| ADCY1 | NM_021116.4 | ATGGATAGCACAGGGGTCCA<br>TTTGGGAGCCGTTTCCATCA | 169 |
| ADCY3 | NM_001320613.2 | CGACCAGAAGGAACGTGTCTA<br>CTCCTCTGTGTAGAAGTCAGCA | 193 |
| ADCY5 | NM_183357.3 | TTTGCTGACATCGAGGGCTT<br>GCCCCGAGACGCAGTAATAA | 163 |
| ADCY8 | NM_001115.3 | AGTGGGGACATTTGTGTGCT<br>CCAGGTTCACAGTTTTGCCC | 186 |
| RAPGEF3 | NM_001098531.4 | CGGAGGGGACACTACTCAAC<br>GGTCTCGGATGAGGTTTGGG | 249 |
| RAPGEF4 | NM_007023.4 | GGATCGCCTGCCTGGATAAA<br>GCCAGGACAGCATACCAGTT | 211 |
| GAPDH | NM_002046.7 | GAAGGTGAAGGTCGGAGTCAA<br>GACAAGCTTCCCGTTCTCAG | 199 |
| 18S rRNA | X03205.1 | TCAACTTTCGATGGTAGTCGCCGT<br>TCCTTGGATGTGGTAGCCGTTTCT | 108 |
